# Deviations from normative functioning underlying emotional episodic memory revealed cross-scale neurodiverse alterations linked to affective symptoms in distinct psychiatric disorders

**DOI:** 10.1101/2024.06.22.600146

**Authors:** Yang Xiao, Mingzhu Li, Xiao Zhang, Yuyanan Zhang, Yuqi Ge, Zhe Lu, Mengying Ma, Yuqing Song, Hao-Yang Tan, Dai Zhang, Weihua Yue, Hao Yan

## Abstract

**Background:** Affective symptoms are a prevalent psychopathological feature in various psychiatric disorders. However, the underlying neurobiological mechanisms are complex and not yet fully understood.

**Methods:** We used normative modelling to establish a reference for neurofunctional activation of functional magnetic resonance imaging based on an emotional episodic memory task, which is frequently used to study affective symptoms in psychiatric disorders. This normative reference was derived from a large dataset of healthy individuals (n = 409), and used to evaluate individualized functional alterations by calculating deviations from this reference in a clinical dataset of 328 participants, which included 168 healthy controls and patients with major depressive disorder (MDD, n = 56), bipolar disorder (BD, n = 31), and schizophrenia (SZ, n = 73). The neurofunctional deviations were mapped to emotional networks with specific emotional functions and used to predict affective symptoms in different mental disorders. The microscale cellular signatures underlying macroscale variations were identified using imaging transcriptomic analysis, and associated with affective symptoms.

**Results:** We observed distinct patterns of cross-scale neural alterations linked to affective symptoms in three psychiatric disorders. Macroscale neural dysfunctions in distinct disorders were embedded into non-overlapping emotional networks and significantly associated with affective symptoms. The oligodendrocytes may mediate the network-specific impairments, and microglia for MDD, astrocytes for BD, and excitatory neurons for SZ as replicable cell-type correlates of affective symptoms.

**Conclusions:** These findings have potential implications for the understanding of unique neuropathological patterns of affective symptoms in distinct psychiatric disorders and improving individualized treatment response.

## Introduction

Affective symptoms, a basic dimension of psychopathology, are highly prevalent across multiple psychiatric conditions (1). In clinic, affective symptoms in psychiatric patients manifest disturbances in emotional processing, the result of which is emotional extremes in perception (2), attention (3), and response (4), and display significant variability in distinct forms of clinical phenotype (5, 6). Clarifying the neurobiological underpinnings related to psychiatric symptoms and behaviors is a major challenge for contemporary research in psychiatry and would help to advance the intervention optimization of psychiatric disorders (7–9). However, the neurobiological alterations underlying affective symptoms in psychiatry are complex and remain poorly understood.

Functional magnetic resonance imaging (fMRI) studies have been a major step forward in characterizing macroscale neural dysfunctions related to affective symptoms in psychiatric patients (5, 10, 11). The emotional episodic memory task (12–14), a widely used paradigm, effectively captures neurofunctional activation associated with adaptive memory-enhancing effects induced by emotional stimuli (15–17) and maladaptive alterations observed in various mental illnesses (18, 19). Disrupted emotional memory retrieval has been reported that may manifest in multiple processes of pathological emotion in psychiatric patients, involving disturbances in emotional attention (20), perceptions (21) and regulation (22), relating to the vulnerability and development of specific clinical affective syndromes (23, 24). Studies using this paradigm have identified altered neurofunctions related to affective symptoms in the amygdala and medial temporal lobe (MTL) system across various mental illnesses (25–27), while also suggesting heterogeneous patterns of neural dysfunctions in distinct diagnoses (28–31).

However, inconsistent results have often been reported, both within one specific diagnosis and across different diagnoses. Increased activity in the amygdala and orbitofrontal gyrus has often been associated with emotional bias in major depressive disorder (MDD) (29, 32), but another study has shown no abnormalities (30). Hypoactivation in the amygdala and occipital gyrus has been recognized in bipolar disorder (BD) compared with MDD patients (33, 34), although an earlier study reported increased activity (35). In individuals diagnosed with schizophrenia (SZ), both hyperactivation and hypoactivation in the prefrontal and parietal cortices have been found (36–38), accompanied by increased or decreased co-activation with the MTL areas (36, 37, 39). In addition, a study has reported that the activation of cingulate cortex was significantly higher in patients with BD relative to SZ (31), but similar tasks found lower activation of cingulate cortex in BD (40). Although these mixed findings could partly be due to differences from task design, one more major reason is the individual differences within and across clinical cohorts, which are often failed to capture by traditional case-control designs (41, 42). An approach that identifies individual-specific alterations and thereby disentangling intersubject heterogeneity of neural dysfunctions that underpin affective symptoms is therefore needed. Normative modelling quantifies each patient as an extreme deviation from normative expectations for MRI-derived phenotypes based on clinical or cognitive characteristics, could avoid confounds arising from intersubject differences to some extent (43, 44).

Complex clinical phenotypes of affective symptoms are not only associated with macroscale functional abnormalities but are also tied to neurobiological alterations at the microscale genetic and cellular levels (45–47). Analyses of postmortem brain samples and animal models have consistently pointed towards cellular processes in the central nervous system, including pathological alterations of neurons and neuroglial cells such as astrocytes and microglia, as an important factor in the development of affective symptoms (48–50). However, due to limitations in spatial resolution and coarse diagnostic classifications, the relationship between microscopic vulnerabilities, macroscale neural dysfunctions, and clinical phenotypes remains poorly understood. The brain-wide gene expression atlas has made it possible to bridge the gap between cross-scale neurofunctional alterations (51) and, by combining with normative modelling framework, could identify the personalized multiscale associations of neural vulnerability.

In the present study, we aimed to investigate multiscale patterns of neural dysfunctions related to affective symptoms in patients with MDD, BD and SZ by considering neurofunctional activation during emotional memory retrieval (Figure 1). By employing two task-fMRI datasets and a normative modelling framework, robust and individualized neurofunctional deviation during emotional memory retrieval was firstly identified. Next, we embedded deviation patterns into macroscale emotional networks to explore the functional annotations of emotional profiling associated with neural variability in distinct disorders, and applied machine-learning approaches to uncover the associations between macroscale alterations and specific effective symptoms dimension in each diagnostic condition. Finally, we conducted transcriptomic association analysis to plausibly determine the cellular abnormalities linked to affective symptoms by integrating macroscale dysfunctions with microscale alterations from the perspective of brain-wide transcriptome and brain single-cell sequencing.

**Figure 1.**
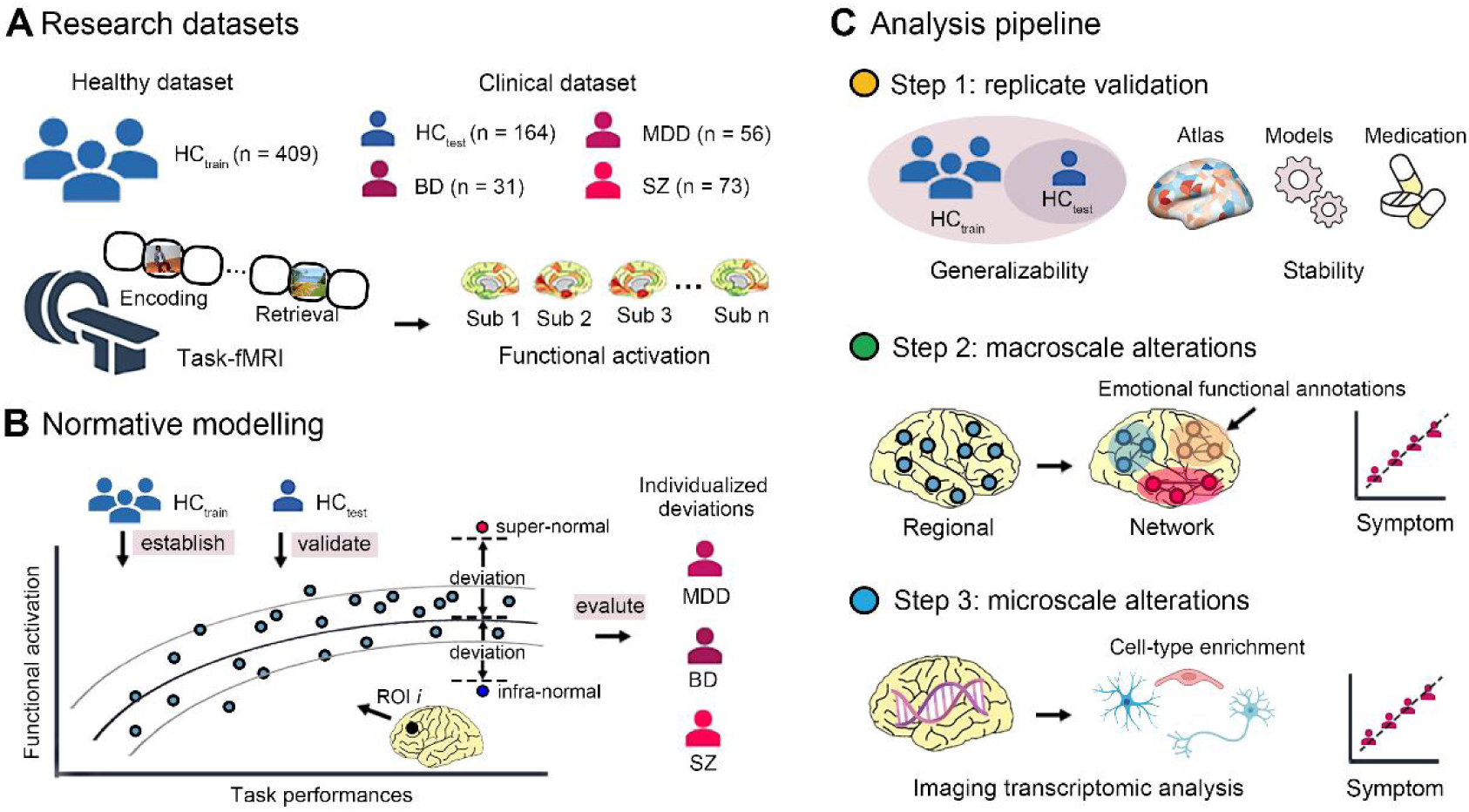
The workflow of the current study design. **(A)** Functional activation for each participant under task-fMRI of emotional episodic memory were extracted from two research datasets (Healthy dataset and Clinical dataset). **(B)** The normative model was first established based on functional activation of each region as response variables and task performances as predicted covariates, drawing from the HC_train_ of Healthy dataset. The HC_test_ in Clinical dataset were further put into normative model for replicate validation, and Individualized alterations were evaluated as deviations relative to normative range at any given brain region at each patient in Clinical dataset. **(C)** Overview of analysis pipeline comprised of three steps. The model generalizability was assessed based on the similarity between the HC_train_ and HC_test_, and the stability of measurements derived from the normative reference was evaluated through replicate validation using different brain parcellations atlas and modeling methods, and validated whether could be influenced by the medication effect (step 1). The macroscale alterations were quantified as the differences between the proportion of individuals showing an extreme deviation of each case and healthy controls at regional level, and further embedded into emotional network for specific functional annotations of emotional profiling. Individual deviations with significant difference were used to predict affective symptom of each psychiatric disorder (step 2). Transcriptomic analysis and cell-type enrichment were further applied to examine the microscale alterations related to macroscale variability, and linked to affective symptom in distinct disorders (step 3). HC, healthy controls; MDD, major depressive disorder; BD, bipolar disorders; SZ, schizophrenia; fMRI, functional magnetic resonance imaging; ROI, region of interest

## Methods and Materials

### 2.1 Participants

The participants of this study were drawn from two datasets. To construct the normative models, Healthy dataset included 409 healthy controls (HC_train_) recruited from the Center for MRI Research, Peking University. The Clinical dataset consisted of 328 individuals: 56 with MDD, 31 with BD, 73 with SZ, and 164 healthy controls (HC_test_). These participants were recruited from the Center for Neuroimaging, Peking University Sixth Hospital. The study was approved by the medical ethics committee of Peking University Sixth Hospital (see Supplement for details of the Ethics Reviews) and all participants provided written informed consent before taking part in the study. All psychiatric patients were assessed and diagnosed using the Diagnostic and Statistical Manual of Mental Disorders, Fourth Edition (DSM-IV) criteria by experienced psychiatric physicians. The severity of depression in MDD patients was accessed using the Hamilton Depression Scale (HAMD), while the clinical symptoms of BD were assessed using both HAMD and the Young Mania Rating Scale (YMRS). The symptoms of patients with SZ were evaluated using the Positive and Negative Syndrome Scale (PANSS). The healthy controls had no current or lifetime history of axis I psychiatric disorders or neurological disorders. See Supplement for additional criteria for participants inclusion and exclusion and quality control details.

### 2.2 Task paradigm

The fMRI paradigm of emotional episodic memory was adopted in Healthy and Clinical dataset, which has been used in previous studies (52), conducting an emotional pictures-encoding and retrieval task during scanning. Briefly, during encoding phase, participants firstly were instructed to determine whether each aversive or neutral picture represented an ‘indoor’ or ‘outdoor’ scene. All participants were not informed about the subsequent retrieval phase before scanning and thus did not realize that they were engaged in a memory task. About 2 minutes after the encoding phase, participants then need to memorize whether the image presented was seen during the encoding session in the case of mixing with half new picture. Finally, both reaction time (RT) and accuracy (ACC) in encoding and retrieval phase were extracted from log files of task performance after scanning. The schematic diagram of the task was shown in Figure 2A and detailed description is in the Supplement.

**Figure 2.**
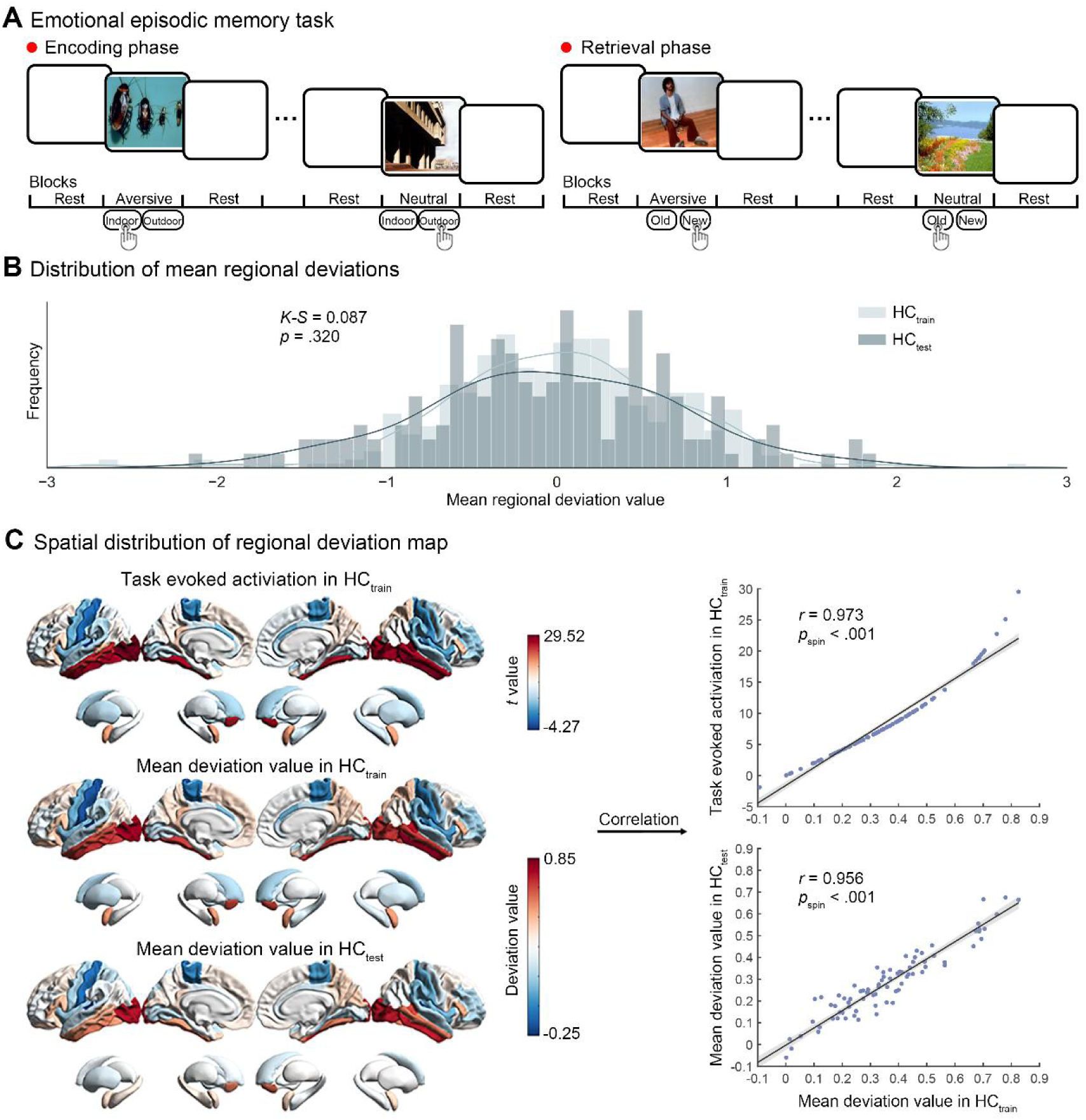
The emotional episodic memory task and generalizability evaluation of normative model. **(A)** The emotional episodic memory task consisted of encoding and retrieval phases and each phase included neutral and aversive scenes. The participants were instructed to determine whether each image represented an ‘indoor’ or ‘outdoor’ scene in encoding phase (left) and respond ‘new’ or for ‘old’ in retrieval phase (right). **(B)** The distribution of mean regional deviations in HC_train_ and HC_test_. There was no significant difference between these two distributions. **(C)** The spatial distribution of regional deviation map in HC_train_ exhibited significant spatial correlations with group-averaged activation and regional deviation map in HC_test_. HC, healthy controls

### 2.3 Image acquisition and preprocessing

Images data of all participants were acquired on a 3.0 T GE Discovery MR750 scanner in the Center for MRI Research, Peking University (Healthy dataset), and the Center for Neuroimaging, Peking University Sixth Hospital (Clinical dataset). For each participant, the high-resolution structural T1-weighted MRI and fMRI blood oxygenation level-dependent (BOLD) signal were acquired, and was preprocessed using the SPM12 (Statistical Parametric Mapping, http://www.fil.ion.ucl.ac.uk/spm) with a standardized protocol whose details can be found elsewhere (53). The details of imaging acquisition and preprocessing can be found in the Supplement.

### 2.4 Functional activation analysis

The analysis of functional activation under emotional episodic memory was based on SPM12 and custom code in MATLAB (The MathWorks, Natick, MA, USA). For each participant, general linear modelling (GLM) analysis was used to identify functions activated by task of each brain regions defined by the Desikan-Killiany (DK) atlas (54). The brain parcellation included 68 cortical regions and 14 subcortical regions, which limited computational burden. Using GLM, regressors were firstly constructed from the rest, aversive and neutral blocks which were then convolved with a canonical double-gamma haemodynamic response function and combined with the temporal derivatives of each main regressor. Then, a first-order autoregressive model was used to remove serial correlations, and ratio normalized to the whole-brain global mean to control for systematic differences in global activity, and temporally filtered using a high-pass filter of 128s to remove low-frequency signal. Next, the contrasts of interest were brain activation responses under rest, aversive and neutral task conditions with the six parameters of head motion as nuisance covariates. Finally, condition-specific regional responses were obtained in three task conditions during retrieval phase, and the “aversive − neutral” activation difference maps in retrieval phase were generated to indicate the effect induced by emotion and, meanwhile, avoid the interference of relevant cognitive factors.

### 2.5 Normative modelling and estimating individual deviations

The normative modelling approach sought to construct a normative distribution to represent the reference range of interest characteristics by calculating the mean and variance of a response variable from a set of predictive factors, thus allowing to characterize a personalized deviation relative to the normative reference (41, 43). We employed gaussian process regression to estimate normative models reflecting relevant neurofunctional activation underlying emotional episodic memory using Predictive Clinical Neuroscience toolkit software (https://pcntoolkit.readthedocs.io/en/latest). Firstly, the activation maps of “aversive − neutral” during retrieval were used as response variables, and predictive factors were comprised of task performances including RT in encoding and retrieval phase and ACC in retrieval phase. Then, the HC_train_ of Healthy dataset was set as training sample to establish the normative reference, the testing sample comprised of the Clinical dataset, including HC_test_, cases with MDD, with BD, and with SZ, was positioned on the normative percentile charts to evaluate individualized deviations for replicate validation and subsequent analyses. We derived a *z* value from model that quantified the deviation from the normative range for each participant in any given brain regions, which can be defined as a measurement of individualized alterations. Finally, to demonstrate the robustness of our main findings, we repeatedly validated our results in the model generalizability and measurements stability. The model generalizability was assessed based on internal validation using cross-validation and external verification based on a testing dataset. Additionally, the stability of individual deviations was validated by considering several potential confounding factors, including different strategies of brain parcellation and modeling method, as well as influences derived from medication effect. Details were described in the Supplement.

### 2.6 Group convergent effect of individual deviations

To quantify the convergent effect of individual neurofunctional deviations in distinct forms of psychiatric disorders, we further defined extreme deviations underlying functional activations by thresholding the individualized deviation (*z* value) maps derived from normative modelling using *z* = ± 2.6 (corresponding to *p* < .005 in standard normal distribution), as was performed in previous studies (41, 55). A group-specific percentage map quantifying the proportion of individuals showing an extreme deviation was calculated, separately for positive and negative extreme deviations. The group convergent effect was assessed as proportion difference of individuals with extreme deviations (Δ percentage map) between each diagnostic group with the HC_test_ of Clinical dataset, aiming at group differences after removing intersubject heterogeneity rather than traditional group-averaged differences. To avoid reliance on a single threshold, the Δ percentage map of each group was estimated based on a threshold-weighted calculation for defining extreme deviations, which integrated results across a range of thresholds (Supplement).

A nonparametric group-based permutation test was conducted to identify significant brain regions of convergent effect at each diagnostic group. Briefly, the percentage maps showing extreme regional deviation in HC_test_ were subtracted from each case’s percentage map to obtain the Δ percentage maps of each disorder at any given brain region. Subsequently, group labels of case group and HC group were shuffled, Δ percentage maps were re-calculated based on permuted group labels. The *p* values were obtained as the proportion of null values that exceeded the observed difference in 10,000 times permutations. and statistical significance was set at 0.05 with multiple comparisons correction based on the false discovery rate (*p* < .05, FDR corrected). The statistically significant differences of positive and negative extreme deviations were respectively defined as supra-normal and infra-normal in current study. The higher or lower infra-normal deviations implied that the hyper- and hypo-inhibition of neural activity, while higher or lower supra-normal deviation suggested hyper- and hypo-excitation, respectively.

Group convergent effect was also assessed at network-level to explore the relevant emotional functional disruptions in distinct disorders, by embedding extreme deviations of brain regions were into large-scale emotional network with specific emotional profiling. A well-recognized framework of four large-scale emotional networks (ENs) with distinct spatial distributions and functional profiles ranging from emotional generation to regulation. (EN 1: working memory and response inhibition in emotional regulation; EN 2: cognitive appraisal and language processing of emotional stimuli; EN 3: emotion perception and generation; EN 4: emotional reactivity), was revealed by a recent meta-analysis study (56), and positioned on DK atlas (Figure S1 and Table S1). Network percentage map was estimated by assigning each region with extreme deviation to one of four large-scale emotional networks, and calculating the proportion of individuals that showed at least one positive and negative extreme deviation in a region within the network. The significant network differences were then evaluated by computing differences between network percentage (Δ network percentage map) of HC_test_ and each diagnostic group using 10,000 times group-based permutation test (*p* < .05, FDR corrected), as at regional level. The details of method can be found in the Supplement and Figure S2, and previous studies (55).

### 2.7 Association with clinical affective symptoms

The machine-learning approaches based on multiple linear regression were conducted to investigate the underlying relationships between individualized neurofunctional deviations and clinical affective symptoms in different diagnostic groups. The affective symptoms in MDD patients were defined using anxiety/somatization factor derived from six-factor domains of HAMD. Considering the lack of specific item scores, the total scores of HAMD were alternatively used as affective symptoms in BD patients. For SZ patients, since a specific assessment scale for depressive symptoms was not adopted, the anxiety/depression factor from five-factor scores of PANSS was used. We used individual deviation of brain regions with significant infra-normal and supra-normal deviation to predict clinical scores in distinct diagnostic category. The nested leave-one-out cross-validation (LOOCV) was adopted to avoid selection biases and evaluate the performance of the linear regression models. The Pearson correlation and root mean square error (RMSE) between the observed scores and predicted scores were used to assess the performance of prediction model. The statistical significance of correlation coefficient was estimated by applying a non-parametric permutation test, which calculated by randomly shuffling the clinical scores and re-calculated Pearson correlation coefficient. An empirical distribution of correlation coefficient was then obtained based on 10,000 times permutation and the significance level threshold was set at *p*_perm_ < .05. Furthermore, we used virtual lesion analysis to determine the important regions and networks in the clinical prediction model by re-calculating the prediction accuracy (correlation coefficient between observed scores and predicted scores) after removing one brain region at a time (i.e., virtual lesion). If the prediction accuracy of model decreased after removing a region, we considered the region to be important for prediction.

### 2.8 Imaging transcriptomic analysis and cell-type enrichment

The imaging transcriptomic analysis was performed to investigate underlying microscale alterations of macroscale dysfunctions linked to affective symptoms in distinct mental illnesses. The brain-wide transcriptome was acquired from post-mortem brain tissue samples of six neurotypical adult donors derived from the Allen Human Brain Atlas (AHBA, http://human.brain-map.org), and preprocessed using abagen toolbox (https://www.github.com/netneurolab/abagen) following established protocols (57), including probe-to-gene reannotation, background noise filtering, sample spatial assignment and normalization. More details of preprocessing of gene expression data can be found in the Supplement. Considering right hemisphere data was available from only two donors, the analysis was restricted to left hemisphere. In total, the preprocessing resulted in a gene expression map (41 brain regions × 15,633 genes) for subsequent analyses. The spatial correlation (Spearman’s ρ) between the expression value of each gene in AHBA and neuroimaging phenotypes were then calculated separately at the group- and individual-level.

To conduct the cell-type enrichment analysis, two cortical single-cell gene sequencing datasets were acquired from previous studies, including the Lake cell-type gene markers obtained from transcriptional data in the frontal and visual cortex (58), as well as the Li cell-type gene markers based on transcriptomic and epigenomic data covering 16 brain regions ranging from prefrontal cortex to cerebellum (59). Following a previous study that conducted a clustering analysis on the above two datasets (60), seven specific cell types were adopted in the current study, including astrocytes, endothelial cells, excitatory neurons, inhibitory neurons, microglia, oligodendrocytes and oligodendrocyte precursors (OPC). As shown in Figure 4A, based on two cell-type gene markers, the FGSEA method was applied to the sorted gene rank calculated from spatial correlation analysis to perform cell-type enrichment analysis (61), and then obtained normalized enrichment scores (NES) at group- and individual-level, respectively. The group-level cell-type enrichment was separately computed at infra- and supra-normal Δ percentage maps at each diagnostic group to capture microscale basis of group convergent neurofunctional alterations, and individualized deviation map was used to calculate individual cell-type enrichment score, as an indicator of cell type-specific molecular contributions underlying macroscale abnormalities. The Pearson correlation was further used to evaluate the relationship between individual cell-type enrichment score and clinical affective symptoms in distinct psychiatric disorders to plausibly determine the cellular abnormalities linked to affective symptoms. All statistical significance was set at 0.05 with multiple comparisons correction by FDR.

## Results

### 3.1 Demographic, clinical characteristics and task performances

The demographical and clinical characteristics are presented in Table 1. A total of 409 HC_train_ participants (mean age, 24.6 ± 3.8 years; age range, 18 to 43 years; 206 female) were recruited in the Healthy dataset. In the Clinical dataset, there were no significant differences in sex (*χ*^2^ = 0.333, *p* = .954) and age (*F* = 2.195, *p* = .089) among the HC_test_, MDD, BD and SZ groups. The statistical analysis of task performances found that emotional memory-enhancing effect was observed across all groups of two datasets, showing significantly greater retrieval accuracy for aversive than for neutral scenes (Table S2). Detailed results of task performances are provided in the Supplement.

**Table 1.**
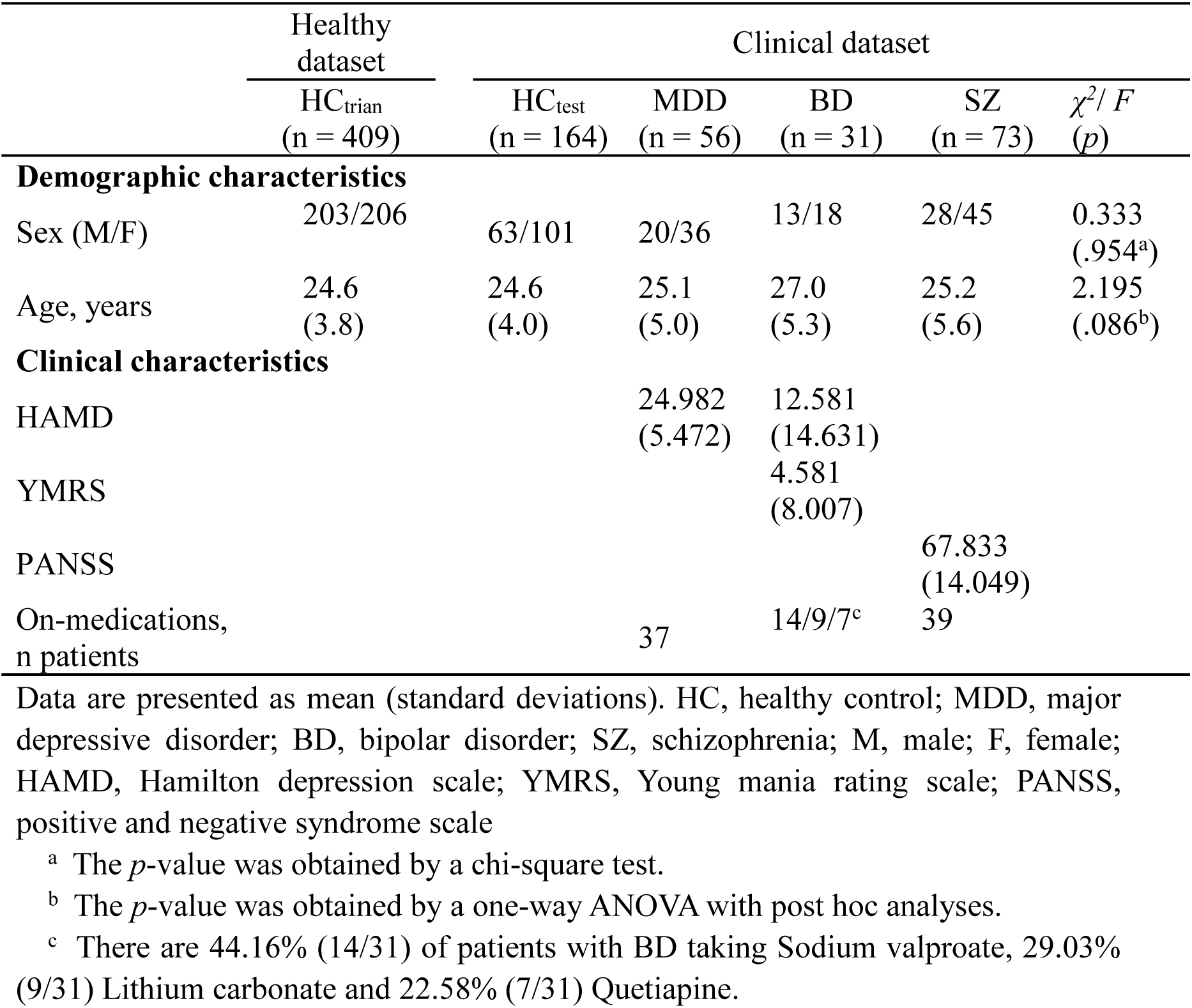
Demographic and clinical characteristics Healthy.

### 3.2 Normative model of neurofunctional activations underlying emotional episodic memory

We constructed a verifiable normative model based on functional activation evoked by the emotional episodic memory paradigm, which was fitted in each region respectively. As shown in Figure 2B, our normative models predicted a normative reference for neurofunctional activations given behavioural performances under emotional episodic memory task, and fitted well in healthy individuals, showing that 99.51% of individuals in HC_train_ and 100% of individuals in HC_test_ were located within the normative range (−2.6 < *z* < 2.6) for the mean regional deviations. There was no significant difference of the distribution of mean regional normative *z* scores between HC_train_ and HC_test_ (Kolmogorov-Smirnov test, *K-S* = 0.087, *p* = .320). Based on general linear model, traditional group-averaged functional activation in HC_train_ showed that regions with higher activation located mostly in the occipital cortices, amygdala and other MTL areas, while those with decreased deviations were mainly in precentral and postcentral gyrus, and insula (*p* < .05, FDR corrected; Figure 2C), consistent with previous studies (62, 63). This spatial distribution of neural activation is highly correlated with individual-level mean distribution patterns of HC_train_ (*r* = 0.973, *p*_spin_ < .001, 10,000 times permutation tests with spatial autocorrelation), indicating that spatial alignment between group-averaged effect and individualized components by characterizing neural activation in given task performances under emotion memory. More importantly, our normative model exhibited a higher generalizability indicated by internal cross-validation (Supplement; Figure S3) and independent external sample validation, showing mean regional deviation map in HC_train_ and HC_test_ was highly associated (Figure 2C, *r* = 0.956, *p*_spin_ < .001, 10,000 times permutation tests with spatial autocorrelation). The normative models established based on behavior performances, sex and age also showed high correlation with current models, which implied that sex and age had no significant effect on our normative model (Supplement; Figure S4).

### 3.3 Clinical heterogeneity of neurofunctional activations underlying emotional episodic memory

Similar to the evaluation of model generalizability in the validation sample of HC_test_ in the Clinical dataset, each patient with specific psychiatric disorder was positioned on the normative percentile charts to calculate individual deviations in each brain region. The results of measurement stability found that individual deviations of all disorders in main findings have high similarity with using different atlas for brain parcellation (Figure S5), and are unrelated to dose of medicine (Figure S6). Considerable intersubject heterogeneity in patients with the same diagnosis was revealed by individualized measurements from normative modelling. About 50% of patients showed extreme deviations in at least one brain region and distributed diffusely in ∼75% regions (Supplement; Figure S7). More specifically, patients with MDD exhibited a higher percentage of supra-normal deviations (90.24%) compared to infra-normal deviations (40.24%). In contrast, BD patients had fewer regions with positive extrema deviations, showing 37.80% supra-normal regions and 65.85% infra-normal regions. For patients with SZ, both positive and negative out-of-range alterations were widely distributed across brain regions, with 93.90% showing supra-normal deviations and 89.02% showing infra-normal deviations (Supplement). As a control analysis, we found that the classical group-level comparison did not reveal any significant regional case-control differences of functional activation (Figure S8). These findings underscored the efficiency of normative modelling in parsing inter-individual differences.

### 3.4 Macroscale alterations linked to affective symptoms

The group-based permutation test was then used to identify convergent effect of individual deviations at each diagnostic group by comparing the proportion of individuals showing significant extreme deviation between HC_test_ and each diagnostic group in the Clinical dataset (Figure 3A). This was done by assessing the observed difference for each disorder in any given brain region (Δ percentage map) against an empirical null distribution. The results showed that there are few spatial overlaps, including as amygdala, insula and supramarginal gyrus, across all mental disorders (*p* < .05, FDR corrected; Figure S9 and Table S3), but the overall spatial distribution is extraordinary different. The infra-normal regions in the patients with MDD mainly located in frontal lobe and temporal lobe, while supra-normal regions were mainly located in superior frontal gyrus, superior temporal gyrus and amygdala (all *p* < .0167 (.05/3), FDR corrected; Figure 3B). The regions with statistically significant difference in MDD patients were summarized in Table S4. Relative to controls, the patients with BD showed infra-normal regions in the occipital lobe, precuneus, hippocampus and amygdala but supra-normal regions in the insula, striatum, and amygdala (all *p* < .0167 (.05/3), FDR corrected; Figure 3E; Table S5). The between-group comparison between SZ patients and HC_test_ showed that the infra-normal regions in insula, hippocampus, and caudate (*p* < .0167 (.05/3), FDR corrected) and supra-normal regions in insula, fusiform gyrus, putamen, and amygdala (*p* < .0167 (.05/3), FDR corrected) in SZ group (Figures 3H; Table S6). These finding were similar with the Δ percentage map using different threshold setting that avoids reliance on a single threshold (Figure S10).

**Figure 3.**
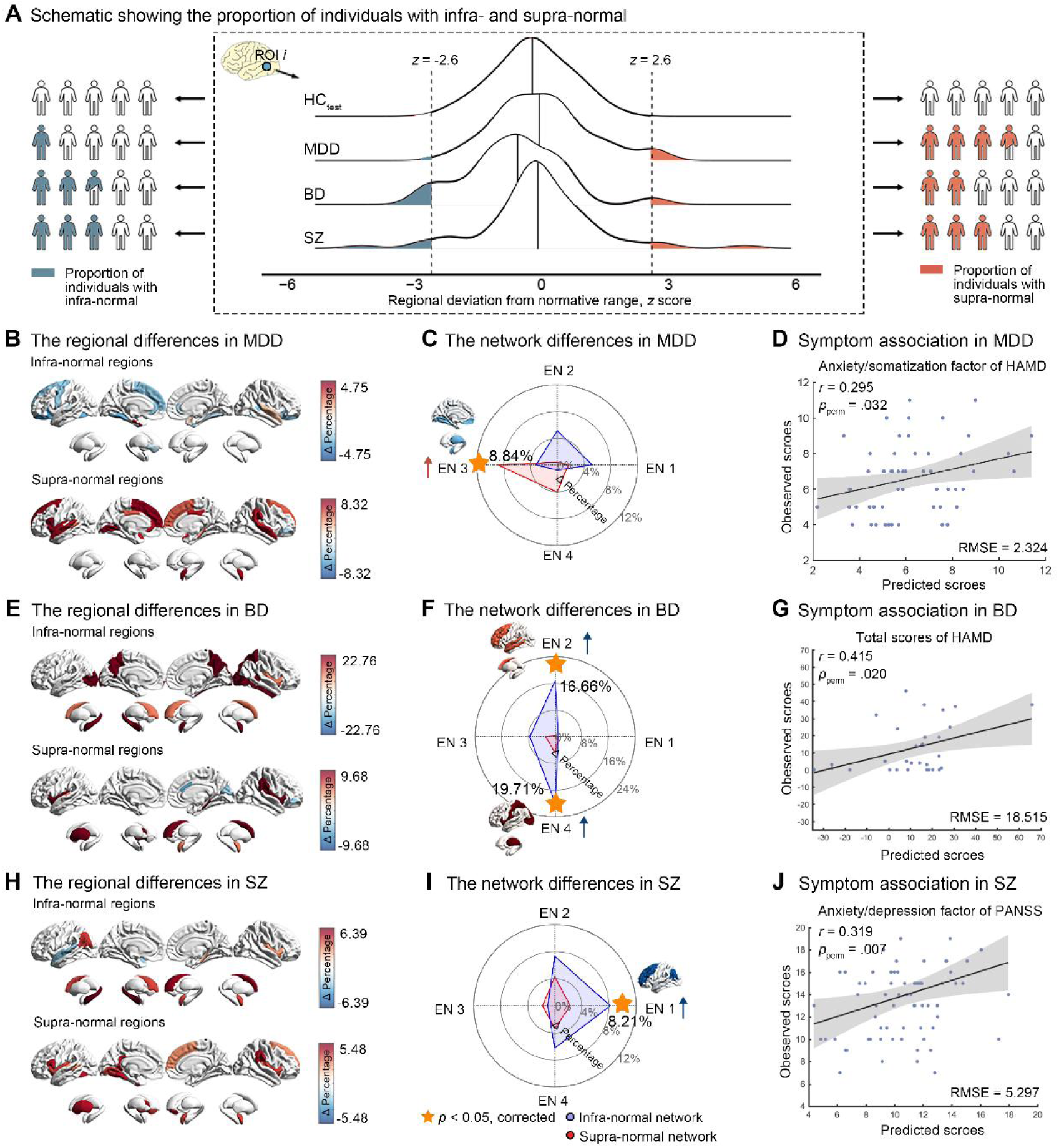
Regional differences and clinical symptom association in MDD, BD and SZ patients. **(A)** Schematic showing the proportion of individuals with infra-normal or supra-normal at any given regions in HC_test_ and each diagnostic group. The infra-normal and supra-normal regions in MDD **(B)**, BD **(E)** and SZ **(H)** patients were identified using 10,000 times group-base permutation test and statistical significance was adjusted by FDR. Regional differences were further enriched into four large-scale emotional networks. The infra-normal and supra-normal alterations at network level in the patients with MDD **(C)**, BD **(F)** and SZ **(I)** were shown. The statistically significance was determined by using 10,000 times group-base permutation test and adjusted by FDR. The scatter plot showed the significant associations between individual neurofunctional deviations and clinical affective symptoms in MDD patients **(D)**, BD **(G)** and SZ **(J)** patients. MDD, major depressive disorder; BD, bipolar disorders; SZ, schizophrenia; HC, healthy controls; EN, emotional network; RMSE, root mean square error; HAMD, Hamilton depression scale; PANSS, positive and negative syndrome scale

Group convergent effect at network level further found that heterogeneous spatial patterns of neural dysfunctions in distinct diagnoses could be attributable to total deviation burden driven by functional alterations of large-scale emotional networks. These networks also covered most of the regions that showed significant functional activation under task conditions (Supplement; Figure S11). Compared to HC, patients with MDD exhibited supra-normal deviations in EN 3 (Δ percentage = 8.84%, *p* = .013, FDR corrected; Figure 3C), indicating higher functional activation in the brain network associated with emotion perception and generation during the emotional episodic memory task; BD patients exhibited infra-normal deviations in networks for appraisal and language processing (EN 2, Δ percentage = 16.66%, *p* = .009, FDR corrected) and emotional reactivity (EN 4, Δ percentage = 19.71%, *p* = .001, FDR corrected) (Figure 3F); SZ patients showed infra-normal deviations in network involving working memory and response inhibition (EN1, Δ percentage = 8.21%, *p* = .025, FDR corrected; Figure 3I). The results from symptom association analyses showed that there are significant correlations between predicted scores and observed scores of anxiety/somatization factor of HAMD in MDD patients (*r* = 0.295, *p*_perm_ = .032, RMSE = 2.324; Figure 3D), total scores of HAMD in BD patients (*r* = 0.415, *p*_perm_ = .020, RMSE = 18.515; Figure 3G), and anxiety/depression factor of PANSS in SZ patients (*r* = 0.319, *p*_perm_ = .007, RMSE = 5.297; Figure 3J). The analysis of important features assessing by virtual lesion analysis have found that emotional networks specific to heterogenous deviations at each psychiatric disorder exhibited maximum predictive weight respectively: EN 3 for MDD, EN 4 for BD, and EN 1 for SZ (Figure S12).

### 3.5 Microscale alterations linked to affective symptoms

Microscale cellular abnormalities underlying macroscale neurofunctional alterations associated with affective symptoms have been identified in distinct forms of mental illnesses, using imaging transcriptomic analysis and single-cell gene data. The results from microscale level showed that macroscale neural dysfunctions exhibit distinct patterns of enrichment across various cell types, but converged on oligodendrocyte across distinct disorders (Figure 4B). More specifically, the results of group-level cell-type enrichment showed that neurofunctional alterations correlated genes in MDD patients were mainly enriched in the microglia for both infra- and supra-normal group effect, and in the oligodendrocyte for supra-normal deviations. For patients with BD, the significantly enriched cell types were in the astrocyte, endothelial cell and oligodendrocyte for infra-normal deviations, and in the oligodendrocyte for supra-normal deviations. The most significant enriched cells in SZ patients are located in the excitatory neurons for both infra- and supra-normal, and in the oligodendrocyte for infra-normal. It is noted that the oligodendrocyte was the most significant enriched cell type in the macroscale neural alterations that showed significant network deviations in distinct diagnostic groups. The analysis of symptom associations of individual cell-type enrichment found that the individual-specific NES of cell types that have significant enrichment at the group-level exhibited significant association with clinical affective symptoms in each psychiatric disorder. The microglia in MDD patients, astrocytes in BD, and excitatory neurons in SZ emerged as replicable cell-level correlates of clinical affective symptoms across two independent single-cell datasets (Figure 4C).

**Figure 4.**
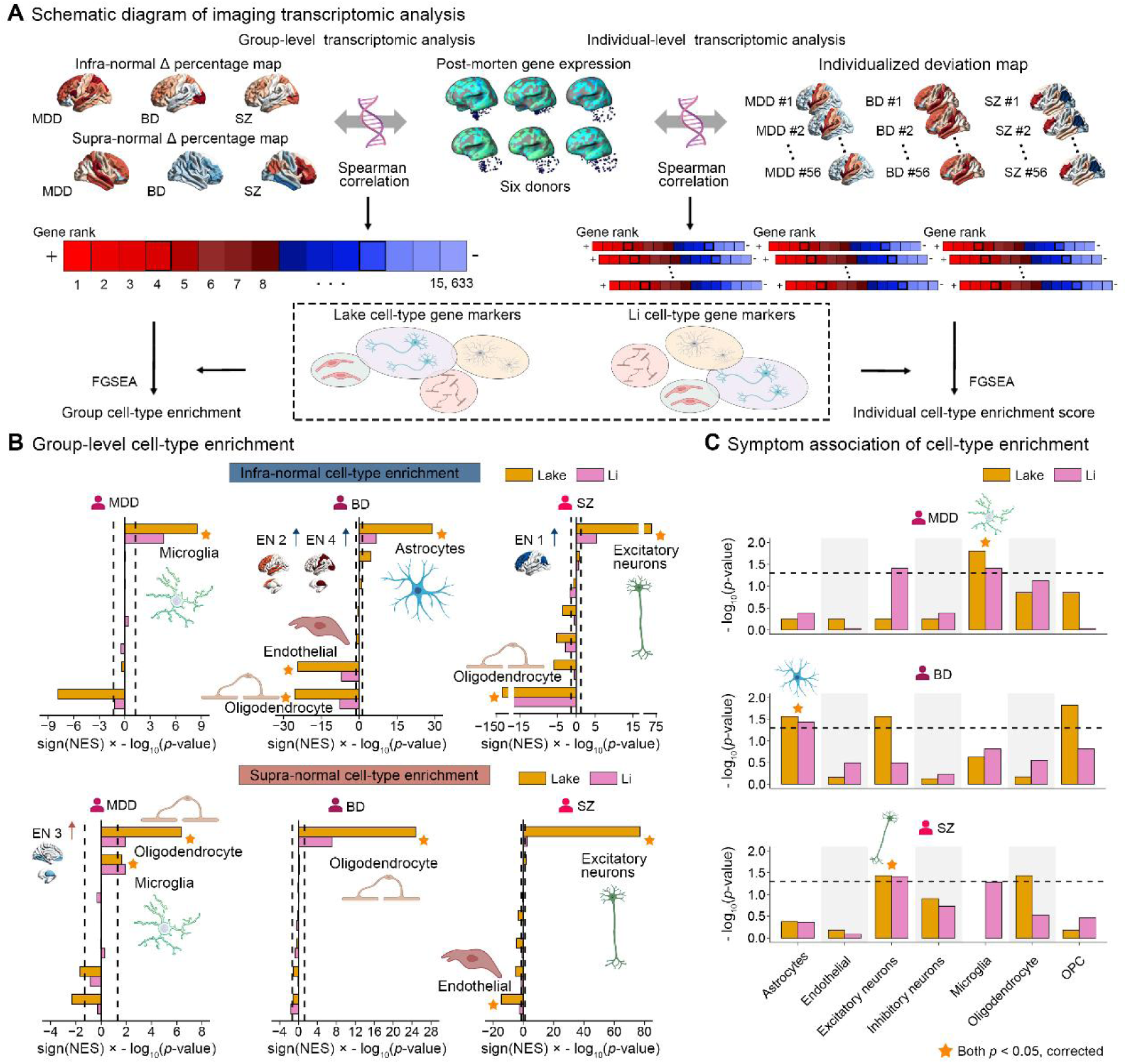
Imaging transcriptomic analysis and cell-type enrichment. **(A)** Schematic diagram of imaging transcriptomic analysis at the group- and individual-level, respectively. Infra- and supra-normal Δ percentage maps at each diagnostic group were spatially correlated with the 15,633 gene expressions of six post-modern brain tissue sample from AHBA using spearman correlation analysis. Individual deviation maps from each patient with psychiatric disorders were also spatially correlated with AHBA gene expressions. Based on two independent cell-type gene makers, the analysis of cell-type enrichment was conducted using FGSEA method in the sorted gene rank to separately calculate group cell-type enrichment and individual-level cell-type enrichment scores. **(B)** Group-level cell-type enrichment. Cell types enriched by infra- and supra-normal deviations were shown across two cell-type gene makers. The horizontal bar plots showed the statistical significance of enrichment that was quantified by sign(NES) × −log10(*p*-value) given by the FGSEA method. The replicable significant cell types across two datasets were labeled. The network group effects of individual deviations corresponding to significant cell-type enrichment were also marked on the side. **(C)** Symptom association of individual cell-type enrichment. The associations between individual-level NES of specific cell type and clinical affective symptoms were examined using the Pearson correlation analysis in distinct psychiatric disorders. The cell types that significantly enriched at the group-level also exhibited significant association with clinical affective symptoms in each psychiatric disorder. The bar plots showed the statistical significance of correlation, and replicable significant cell types across two datasets were labeled. MDD, major depressive disorder; BD, bipolar disorders; SZ, schizophrenia; EN, emotional network; FGSEA, fast gene set enrichment analysis; OPC, oligodendrocyte precursors

## Discussion

The present study aims to identify cross-scale neural alterations that underpin affective symptoms in three major psychiatric disorders. Our findings revealed that macroscale neural dysfunctions associated with specific domains of affective symptoms across different mental illnesses, after removing the confounds from clinical heterogeneity by applying the normative modelling in task-fMRI of emotional memory paradigm. Altered neurofunctions in different disorders could be embedded into non-overlapping emotional networks with specific functional profiling, and the main disruptions of emotional functions related to affective symptoms in distinct disorders are identified. Our results also revealed microscale cellular signatures underlying macroscale functional abnormalities and the cell type-specific processes linked to affective symptoms in various mental illnesses. Overall, our study provides critical insights for better understanding the neural mechanisms underlying specific affective symptoms in distinct psychiatric disorders.

Mapping complex clinical symptoms to neurobiological alterations is significant for exposing more effective interventions for patients with complex psychiatric symptoms (8). Previous studies focusing on the neural activities of affective symptoms commonly applied the emotional episodic memory task to characterizing relevant brain regions, and indicated better performances compared with task-free neuroimaging markers (52, 64). However, limiting methodology and high intersubject heterogeneity, objective and reliable neural correlates related to affective symptoms in multiple psychiatric disorders still have not been revealed. According to the evaluation of sample sizes for traditional case-control designs (Table S7), previous studies have often used a relatively small size of samples. Using normative modelling techniques, we were able to not be limited by the requirements of sample size, and disentangle the clinical heterogeneity to some extent, based on a nonparametric estimation by incorporating neural alterations into a continuous spectrum derived from a large healthy population. Our results from normative modelling also revealed considerable inconsistency in the functional alterations under emotional memory task, showing only a low proportion of the patients shared the consistent extreme deviation for any single brain region (Figure S7). After removing the confounds from clinical heterogeneity, different degrees of neurobiological alterations associated with affective symptom dimension were measured at each patient in distinct diagnostic categories. Association analyses revealed individual deviations significant predicted the affective symptom dimension in specific disorder condition, further determining that results from our paradigm and model might be a promising predictor for clinical affective symptoms. More importantly, strict replicate validations were applied in our model, ranging from model generalizability to measurements stability based on multiple strategies (Figure S3-S6). Among them, medication has long been considered as an important confounding factor, contributing to the lack of consistent findings across studies (65). For instance, the use of psychoactive drugs may impact on neurocognitive systems supporting procedural learning and conditioning and thereby modulate emotional memory processes (63). The stability of the medication effects has critical implications for understanding mechanisms underlying impaired emotional functions across different diagnostic conditions.

To characterize macroscale pattern of neural alterations underlying affective symptoms and better compare with previous studies, the group convergent effects of individual neurofunctional deviations at each diagnostic group were estimated as in a previous study (55). The results showed that the macroscale alterations in distinct disorders are largely distinct but have some overlap. Neural dysfunctions intersected across all mental disorders were found in hyperactivation of the right amygdala, left insula and right supramarginal gyrus (Figure S9). The amygdala has been widely emphasized as underlying emotional impairments in mental illness, engaging abnormal responses to external negative stimuli (5, 66), while the insula is associated with inflexible updating and preferential maintenance of negative or threatening stimuli (67, 68). The co-hyperactivation of amygdala and insula may indicate that excessive attention to negative stimulation may produce an interference effect on memory of stimuli, contributing to systematic distortions in emotional information processing in mental disorders. Additionally, growing evidence suggests that emotional memory bias is not limited to attention and emotion-related problems but is embedded in stable negative self-referential schemas (29, 69). The supramarginal gyrus, part of the default mode network, is linked to self-focused rumination and abnormal self-referential processes (70, 71). Its abnormal activation may drive weakened cognitive structures to produce interpretive biases in mental disorders (14).

The heterogeneous patterns of macroscale functional impairments in different diagnostic groups were embedded into non-overlapping large-scale emotional networks. After embedding to network level, the proportions of individuals with extreme deviations have increased relative to at regional level (Figure S13), showing heterogeneous deficits of distinct disorders could be aggregated within lesioned large-scale emotional networks. Specifically, we found that patients with MDD exhibited hyperactivations in a network related to emotion perception and generation, which comprised of the amygdala, fusiform, and medial orbitofrontal gyrus. Cognitive theories of MDD hold that emotional abnormalities derive from a negative schema for representing negative knowledge and experiences (72), which could alter perceptions and evaluations of external stimuli (73) and its weakening has been hypothesized to underlie recovery (74, 75). A recent study also suggested that regulating emotion perception could be used to reduce emotional memory and improve depressive symptoms (22). SZ patients showed infra-normal deviations in a cognitive emotion regulation network strongly associated with working memory and response inhibition, which is a frontoparietal network consisting of dorsolateral prefrontal cortex and inferior parietal gyrus. The emotional flattening and anhedonia in SZ patients have been reported may contribute to the overuse of suppression strategy, and thereby disrupt emotional appraisal and recognition (76, 77), which align with our findings of dysfunctions of cognitive emotion regulation network. In contrast, individuals with BD were associated with two emotional networks, jointly responsible for integrating emotional reactivity and cognition. This finding provided consistent evidence for support that the effective segregation of neural mechanisms between over-sensitive reactivity and cognitive dysregulation in different emotional states of BD (78), suggesting set state-specific psychotherapeutic targets at the depressive and manic state of BD patients (79). Analyses of symptom associations further found that parallel deficits of large-scale emotional networks in specific psychiatric conditions exhibited maximum predictive weight in the midst of each predictive model (Figure S12). This investigation has critical heuristic value for guiding the optimization of clinical treatment strategies for affective symptoms.

Microscale cellular abnormalities underlying macroscale neurofunctional deviations could further be revealed using imaging transcriptomic analysis and cellular decoding. Considering previous studies indicated that those microscopic cellular alterations possibly induced macroscopic functional variations linked to affective symptoms (45–47), our analyses mainly focused on the cellular basis of functional deviations. The results from group-level cell-type enrichment showed the specific cell types were associated with macroscale alterations in distinct disorders, suggesting heterogeneous cellular abnormalities across distinct mental illnesses, but a convergence of significant network deviations was found in the oligodendrocytes in all disorders. The consistent enrichment in oligodendrocytes may imply that network-level neurofunctional alterations at each diagnostic group may be mediated by dysfunction in integration or segregation within the network, that is, the impaired patterns of functional deviations at the circuit-level. To prove it, using lesion network mapping methodology (8, 80), circuit group convergent effects of individual deviations were calculated based on seed regions with significant extreme deviations, as conducted in a previous study (55). The significant functional circuit was further located into four ENs and found the most significant circuit was focused on specific emotional networks of each case group (Supplement; Figure S14 and Tables S8-10). This raised a hypothesis that the network characteristic with emotional functional profiling were decoded by abnormal connectivity integrations and thereby contributing to clinical affective symptoms. Benefit from measuring individual functional deviations from normative modelling, we could further determine whether the identified cellular characteristics can be used as a potential microscale vulnerability risk factor reflecting clinical symptoms. We found that individual-specific cellular transcriptomic profiles of the identified group-level cell types were significantly associated with affective symptoms. It is suggested that cellular abnormalities play a role in mediating the relationship with macroscale neural activities and external affective symptoms.

In clinic, patients with MDD show persistent low mood and rumination about negative thoughts (81), whereas the hallmark feature of BD patients is fluctuating episodes of depression and mania (82). Diminished emotion expression or affective flattening are also a core feature of persistent functional disability in SZ and may heighten impending psychotic relapse (77, 83). Considering the heterogeneous nature of affective symptoms in different psychiatric disorders, the cross-scale neural alterations comprising of macroscale diagnosis-specific emotional networks and microscale cellular abnormalities may provide enlightening evidences of unique dysfunctional neural characteristics. These findings are consistent with previous theoretical models and research summaries (6, 28), and our research revealed the neurodiverse dysfunctions of affective symptoms in different mental disorders for the first time in empirical research.

Several issues need to be considered. First, the sample size of clinical diagnostic group in the current dataset was relatively small. Although we used a nonparametric group-based permutation test to identify statistically significant deviations in each case group, which is not affected by sample size, a larger sample would help to better parse clinical heterogeneity. Second, data on depressive severity in BD patients was partially missing, making it impossible to calculate the anxiety/somatization factor of HAMD to represent affective symptoms in BD. Additionally, a greater number and more comprehensive assessment will help better characterize the affective symptoms of different mental disorders. Finally, most BD patients in this study were in the euthymic stage. Further study with more patients in different stages is imperative to assess the reproducibility of our findings.

## Conclusion

To summarize, this study proposed cross-scale neural alterations underlying affective symptoms in multiple psychiatric disorders. Based on a stable and cross-sample verifiable normative model of functional activation under emotional episodic memory task, we identified macroscale heterogenous patterns of neurofunctional alterations annotating specific emotional profiling from large-scale emotional network in three common psychiatric disorders. The microscale specific cellular abnormalities mediated the macroscale diagnosis-specific network dysfunctions, and as a potential risk factor of biological vulnerability reflecting clinical symptoms. These findings are a step forward in understanding the cross-scale neurobiology underlying basic dimensions of affectivity and provide a novel insight for translating into clinical management and treatment approaches.

## Acknowledgements

We extend our gratitude to all participants in this study. This study was supported by the National Natural Science Foundation of China (82071505, 81771443, 81361120395), the Clinical Medicine Plus X – Young Scholars Project, Peking University the Fundamental Research Funds for the Central Universities (PKU2024LCXQ046) and Capital’s Funds for Health Improvement and Research (2024-2-4115, 2024-1-4111).

## References

1. Association AP (2013): Diagnostic and statistical manual of mental disorders (5th ed). American Psychiatric Association.

2. Whitton AE, Treadway MT, Pizzagalli DA (2015): Reward processing dysfunction in major depression, bipolar disorder and schizophrenia. Current opinion in psychiatry. 28:7–12.

3. Todd J, Coutts-Bain D, Wilson E, Clarke P (2023): Is attentional bias variability causally implicated in emotional vulnerability? A systematic review and meta-analysis. Neuroscience and biobehavioral reviews. 146:105069.

4. Caballero C, Nook EC, Gee DG (2023): Managing fear and anxiety in development: A framework for understanding the neurodevelopment of emotion regulation capacity and tendency. Neuroscience and biobehavioral reviews. 145:105002.

5. McTeague LM, Rosenberg BM, Lopez JW, Carreon DM, Huemer J, Jiang Y, et al. (2020): Identification of common neural circuit disruptions in emotional processing across psychiatric disorders. The American journal of psychiatry. 177:411–421.

6. Sheppes G, Suri G, Gross JJ (2015): Emotion regulation and psychopathology. Annual review of clinical psychology. 11:379–405.

7. Martino M, Magioncalda P (2024): A working model of neural activity and phenomenal experience in psychosis. Molecular psychiatry. Advance online publication.

8. Fox MD (2018): Mapping symptoms to brain networks with the human connectome. The New England journal of medicine. 379:2237–2245.

9. Martino M, Magioncalda P (2024): A three-dimensional model of neural activity and phenomenal-behavioral patterns. Molecular psychiatry. 29:639–652.

10. de Quervain D, Schwabe L, Roozendaal B (2017): Stress, glucocorticoids and memory: implications for treating fear-related disorders. Nature reviews Neuroscience. 18:7–19.

11. Fastenrath M, Spalek K, Coynel D, Loos E, Milnik A, Egli T, et al. (2022): Human cerebellum and corticocerebellar connections involved in emotional memory enhancement. Proceedings of the National Academy of Sciences of the United States of America. 119:e2204900119.

12. B JMB, Persaud MR, Smith D, Kapczinski FP, Frey BN (2019): Explicit emotional memory biases in mood disorders: A systematic review. Psychiatry research. 278:162–172.

13. Neumann A, Blairy S, Lecompte D, Philippot P (2007): Specificity deficit in the recollection of emotional memories in schizophrenia. Consciousness and cognition. 16:469–484.

14. Lavigne KM, Deng J, Raucher-Chéné D, Hotte-Meunier A, Voyer C, Sarraf L, et al. (2024): Transdiagnostic cognitive biases in psychiatric disorders: A systematic review and network meta-analysis. Progress in neuro-psychopharmacology & biological psychiatry. 129:110894.

15. Hamann S (2001): Cognitive and neural mechanisms of emotional memory. Trends in cognitive sciences. 5:394–400.

16. Faul L, LaBar KS (2023): Mood-congruent memory revisited. Psychological review. 130:1421–1456.

17. Fitzgerald DA, Arnold JF, Becker ES, Speckens AE, Rinck M, Rijpkema M, et al. (2011): How mood challenges emotional memory formation: an fMRI investigation. NeuroImage. 56:1783–1790.

18. LaBar KS, Cabeza RJNRN (2006): Cognitive neuroscience of emotional memory. Nature reviews Neuroscience. 7:54–64.

19. Wolf OT (2008): The influence of stress hormones on emotional memory: relevance for psychopathology. Acta psychologica. 127:513–531.

20. Leppänen JMJCoip (2006): Emotional information processing in mood disorders: a review of behavioral and neuroimaging findings. Current opinion in psychiatry. 19:34–39.

21. Hamilton JP, Gotlib IH (2008): Neural substrates of increased memory sensitivity for negative stimuli in major depression. Biological psychiatry. 63:1155–1162.

22. Hayes BK, Harikumar A, Ferguson LA, Dicker EE, Denny BT, Leal SL (2023): Emotion regulation during encoding reduces negative and enhances neutral mnemonic discrimination in individuals with depressive symptoms. Neurobiology of learning and memory. 205:107824.

23. Haas BW, Canli T (2008): Emotional memory function, personality structure and psychopathology: a neural system approach to the identification of vulnerability markers. Brain research reviews. 58:71–84.

24. Ledoux JE, Muller J (1997): Emotional memory and psychopathology. Philosophical transactions of the Royal Society of London Series B, Biological sciences. 352:1719–1726.

25. Girardeau G, Inema I, Buzsáki G (2017): Reactivations of emotional memory in the hippocampus-amygdala system during sleep. Nature neuroscience. 20:1634–1642.

26. Fan X, Mocchi M, Pascuzzi B, Xiao J, Metzger BA, Mathura RK, et al. (2024): Brain mechanisms underlying the emotion processing bias in treatment-resistant depression. Nature Mental Health. 2:583–592.

27. Roozendaal B, McEwen BS, Chattarji S (2009): Stress, memory and the amygdala. Nature reviews Neuroscience. 10:423–433.

28. Phillips ML, Drevets WC, Rauch SL, Lane R (2003): Neurobiology of emotion perception II: Implications for major psychiatric disorders. Biological psychiatry. 54:515–528.

29. Ghosh VE, Gilboa A (2014): What is a memory schema? A historical perspective on current neuroscience literature. Neuropsychologia. 53:104–114.

30. Rai S, Griffiths K, Breukelaar IA, Barreiros AR, Chen W, Boyce P, et al. (2021): Investigating neural circuits of emotion regulation to distinguish euthymic patients with bipolar disorder and major depressive disorder. Bipolar disorders. 23:284–294.

31. Whalley HC, McKirdy J, Romaniuk L, Sussmann J, Johnstone EC, Wan HI, et al. (2009): Functional imaging of emotional memory in bipolar disorder and schizophrenia. Bipolar disorders. 11:840–856.

32. Ai H, Opmeer EM, Veltman DJ, van der Wee NJ, van Buchem MA, Aleman A, et al. (2015): Brain activation during emotional memory processing associated with subsequent course of depression. Neuropsychopharmacology. 40:2454–2463.

33. Young KD, Bodurka J, Drevets WC (2016): Differential neural correlates of autobiographical memory recall in bipolar and unipolar depression. Bipolar disorders. 18:571–582.

34. Tseng WL, Bones BL, Kayser RR, Olsavsky AK, Fromm SJ, Pine DS, et al. (2015): An fMRI study of emotional face encoding in youth at risk for bipolar disorder. European psychiatry. 30:94–98.

35. Perlman SB, Almeida JR, Kronhaus DM, Versace A, Labarbara EJ, Klein CR, et al. (2012): Amygdala activity and prefrontal cortex-amygdala effective connectivity to emerging emotional faces distinguish remitted and depressed mood states in bipolar disorder. Bipolar disorders. 14:162–174.

36. Hall J, Harris JM, McKirdy JW, Johnstone EC, Lawrie SM (2007): Emotional memory in schizophrenia. Neuropsychologia. 45:1152–1159.

37. Sergerie K, Armony JL, Menear M, Sutton H, Lepage M (2009): Influence of emotional expression on memory recognition bias in schizophrenia as revealed by fMRI. Schizophrenia bulletin. 36:800–810.

38. Kim H (2011): Neural activity that predicts subsequent memory and forgetting: a meta-analysis of 74 fMRI studies. NeuroImage. 54:2446–2461.

39. Rasetti R, Mattay VS, White MG, Sambataro F, Podell JE, Zoltick B, et al. (2014): Altered hippocampal-parahippocampal function during stimulus encoding: a potential indicator of genetic liability for schizophrenia. JAMA psychiatry. 71:236–247.

40. Zhang L, Opmeer EM, Ruhé HG, Aleman A, van der Meer L (2015): Brain activation during self- and other-reflection in bipolar disorder with a history of psychosis: comparison to schizophrenia. NeuroImage Clinical. 8:202–209.

41. Marquand AF, Rezek I, Buitelaar J, Beckmann CF (2016): Understanding heterogeneity in clinical cohorts using normative models: beyond case-control studies. Biological psychiatry. 80:552–561.

42. Rutherford S, Kia SM, Wolfers T, Fraza C, Zabihi M, Dinga R, et al. (2022): The normative modeling framework for computational psychiatry. Nature protocols. 17:1711–1734.

43. Marquand AF, Kia SM, Zabihi M, Wolfers T, Buitelaar JK, Beckmann CF (2019): Conceptualizing mental disorders as deviations from normative functioning. Molecular psychiatry. 24:1415–1424.

44. Haas SS, Ge R, Agartz I, Amminger GP, Andreassen OA, Bachman P, et al. (2024): Normative modeling of brain morphometry in clinical high risk for psychosis. JAMA psychiatry. 81:77–88.

45. Price JL, Drevets WC (2010): Neurocircuitry of mood disorders. Neuropsychopharmacology. 35:192–216.

46. Ramos-Prats A, Matulewicz P, Edenhofer ML, Wang KY, Yeh CW, Fajardo-Serrano A, et al. (2024): Loss of mGlu(5) receptors in somatostatin-expressing neurons alters negative emotional states. Molecular psychiatry. 29:2774–2786.

47. Li H, Namburi P, Olson JM, Borio M, Lemieux ME, Beyeler A, et al. (2022): Neurotensin orchestrates valence assignment in the amygdala. Nature. 608:586–592.

48. Hu X, Zhu Q, Lou T, Hu Q, Li H, Xu Y, et al. (2024): Pan-ErbB inhibition impairs cognition via disrupting myelination and aerobic glycolysis in oligodendrocytes. Proceedings of the National Academy of Sciences of the United States of America. 121:e2405152121.

49. Klawonn AM, Fritz M, Castany S, Pignatelli M, Canal C, Similä F, et al. (2021): Microglial activation elicits a negative affective state through prostaglandin-mediated modulation of striatal neurons. Immunity. 54:225–234.e226.

50. Wahis J, Baudon A, Althammer F, Kerspern D, Goyon S, Hagiwara D, et al. (2021): Astrocytes mediate the effect of oxytocin in the central amygdala on neuronal activity and affective states in rodents. Nature neuroscience. 24:529–541.

51. Hawrylycz MJ, Lein ES, Guillozet-Bongaarts AL, Shen EH, Ng L, Miller JA, et al. (2012): An anatomically comprehensive atlas of the adult human brain transcriptome. Nature. 489:391–399.

52. Chen Q, Ursini G, Romer AL, Knodt AR, Mezeivtch K, Xiao E, et al. (2018): Schizophrenia polygenic risk score predicts mnemonic hippocampal activity. Brain. 141:1218–1228.

53. Acton PD, Friston KJ (1998): Statistical parametric mapping in functional neuroimaging: beyond PET and fMRI activation studies. European journal of nuclear medicine. 25:663–667.

54. Desikan RS, Ségonne F, Fischl B, Quinn BT, Dickerson BC, Blacker D, et al. (2006): An automated labeling system for subdividing the human cerebral cortex on MRI scans into gyral based regions of interest. NeuroImage. 31:968–980.

55. Segal A, Parkes L, Aquino K, Kia SM, Wolfers T, Franke B, et al. (2023): Regional, circuit and network heterogeneity of brain abnormalities in psychiatric disorders. Nature neuroscience. 26:1613–1629.

56. Morawetz C, Riedel MC, Salo T, Berboth S, Eickhoff SB, Laird AR, et al. (2020): Multiple large-scale neural networks underlying emotion regulation. Neuroscience and biobehavioral reviews. 116:382–395.

57. Arnatkeviciute A, Fulcher BD, Fornito A (2019): A practical guide to linking brain-wide gene expression and neuroimaging data. NeuroImage. 189:353–367.

58. Lake BB, Chen S, Sos BC, Fan J, Kaeser GE, Yung YC, et al. (2018): Integrative single-cell analysis of transcriptional and epigenetic states in the human adult brain. Nature biotechnology. 36:70–80.

59. Li M, Santpere G, Imamura Kawasawa Y, Evgrafov OV, Gulden FO, Pochareddy S, et al. (2018): Integrative functional genomic analysis of human brain development and neuropsychiatric risks. Science. 362(6420): eaat7615.

60. Seidlitz J, Nadig A, Liu S, Bethlehem RAI, Vértes PE, Morgan SE, et al. (2020): Transcriptomic and cellular decoding of regional brain vulnerability to neurogenetic disorders. Nature communications. 11:3358.

61. Korotkevich G, Sukhov V, Budin N, Shpak B, Artyomov MN, Sergushichev A (2021): Fast gene set enrichment analysis. Preprint at bioRxiv.

62. Dolcos F, Katsumi Y, Weymar M, Moore M, Tsukiura T, Dolcos S (2017): Emerging directions in emotional episodic memory. Frontiers in psychology. 8:1867.

63. Doss MK, Samaha J, Barrett FS, Griffiths RR, de Wit H, Gallo DA, et al. (2024): Unique effects of sedatives, dissociatives, psychedelics, stimulants, and cannabinoids on episodic memory: A review and reanalysis of acute drug effects on recollection, familiarity, and metamemory. Psychological review. 131:523–562.

64. Dickerson BC, Eichenbaum H (2010): The episodic memory system: neurocircuitry and disorders. Neuropsychopharmacology. 35:86–104.

65. Doss MK, de Wit H, Gallo DA (2023): The acute effects of psychoactive drugs on emotional episodic memory encoding, consolidation, and retrieval: a comprehensive review. Neuroscience and biobehavioral reviews. 150:105188.

66. Janak PH, Tye KM (2015): From circuits to behaviour in the amygdala. Nature. 517:284–292.

67. Yaple ZA, Tolomeo S, Yu R (2021): Mapping working memory-specific dysfunction using a transdiagnostic approach. NeuroImage Clinical. 31:102747.

68. Zhang JN, Xiong KL, Qiu MG, Zhang Y, Xie B, Wang J, et al. (2013): Negative emotional distraction on neural circuits for working memory in patients with posttraumatic stress disorder. Brain research. 1531:94–101.

69. Bogie BJM, Kapczinski FP, McCabe RE, McKinnon MC, Frey BN (2020): Emotional reactivity and explicit emotional memory biases in major depressive disorder during euthymia. Psychiatry research. 285:112847.

70. Chen X, Chen NX, Shen YQ, Li HX, Li L, Lu B, et al. (2020): The subsystem mechanism of default mode network underlying rumination: a reproducible neuroimaging study. NeuroImage. 221:117185.

71. Zhou HX, Chen X, Shen YQ, Li L, Chen NX, Zhu ZC, et al. (2020): Rumination and the default mode network: meta-analysis of brain imaging studies and implications for depression. NeuroImage. 206:116287.

72. Vrijsen JN, Ikani N, Souren P, Rinck M, Tendolkar I, Schene AH (2023): How context, mood, and emotional memory interact in depression: a study in everyday life. Emotion. 23:41–51.

73. Clark DA, Beck AT (2010): Cognitive theory and therapy of anxiety and depression: convergence with neurobiological findings. Trends in cognitive sciences. 14:418–424.

74. Bockting CL, Hollon SD, Jarrett RB, Kuyken W, Dobson K (2015): A lifetime approach to major depressive disorder: The contributions of psychological interventions in preventing relapse and recurrence. Clinical psychology review. 41:16–26.

75. Scheepens DS, van Waarde JA, Ten Doesschate F, Westra M, Kroes MCW, Schene AH, et al. (2020): Effectiveness of emotional memory reactivation vs control memory reactivation before electroconvulsive therapy in adult patients with depressive disorder: a randomized clinical trial. JAMA network open. 3:e2012389.

76. Guimond S, Padani S, Lutz O, Eack S, Thermenos H, Keshavan M (2018): Impaired regulation of emotional distractors during working memory load in schizophrenia. Journal of psychiatric research. 101:14–20.

77. O’Driscoll C, Laing J, Mason O (2014): Cognitive emotion regulation strategies, alexithymia and dissociation in schizophrenia, a review and meta-analysis. Clinical psychology review. 34:482–495.

78. Frodl T (2023): Adaptive and maladaptive emotion regulation in bipolar disorder. Acta psychiatrica Scandinavica. 148:469–471.

79. Oliva V, De Prisco M, Fico G, Possidente C, Fortea L, Montejo L, et al. (2023): Correlation between emotion dysregulation and mood symptoms of bipolar disorder: A systematic review and meta-analysis. Acta psychiatrica Scandinavica. 148:472–490.

80. Boes AD, Prasad S, Liu H, Liu Q, Pascual-Leone A, Caviness VS, Jr., et al. (2015): Network localization of neurological symptoms from focal brain lesions. Brain. 138:3061–3075.

81. Malhi GS, Mann JJ (2018): Depression. Lancet (London, England). 392:2299–2312.

82. Vieta E, Berk M, Schulze TG, Carvalho AF, Suppes T, Calabrese JR, et al. (2018): Bipolar disorders. Nature reviews Disease primers. 4:18008.

83. Zou YM, Ni K, Yang ZY, Li Y, Cai XL, Xie DJ, et al. (2018): Profiling of experiential pleasure, emotional regulation and emotion expression in patients with schizophrenia. Schizophrenia research. 195:396–401.

