## Supplemental Materials for "Deviations from normative functioning underlying emotional episodic memory revealed cross-scale neurodiverse alterations linked to affective symptoms in distinct psychiatric disorders"

**SUPPLEMENTARY MATERIALS**

### Ethics reviews

The authors assert that all procedures contributing to this work comply with the ethical standards of the relevant national and institutional committees on human experimentation and with the [Helsinki Declaration](https://www.wma.net/policies-post/wma-declaration-of-helsinki-ethical-principles-for-medical-research-involving-human-subjects/) of 1975, as revised in 2013. The study was approved by the medical ethics committee of Peking University Sixth Hospital, and approval of corresponding experimental protocols (Ethics Review No. 32 of 2013 for the Healthy dataset and No. 32 of 2015 for BP, No. 32 of 2017 for MDD and No. 2 of 2021 for SZ and HC_test_ in the Clinical dataset).

### The inclusion and exclusion criteria of datasets

The participants of this study were taken from two datasets. The healthy dataset included 409 healthy controls (HC_train­_) were recruited from the Center for MRI Research, Peking University. The clinical dataset consisted of 328 individuals: 56 with major depressive disorder (MDD), 31 with bipolar disorder (BD), 73 with schizophrenia (SZ), and 164 healthy controls (HC_test_). All participants of the clinical dataset were recruited from the Center for Neuroimaging, Peking University Sixth Hospital. All participants were included if they: 1) were Chinese of Han ancestry; 2) age ranging from 18 to 43 years old; 3) were right-handed based on the evaluation of the handedness questionnaire. The healthy samples were excluded if they: 1) had current or history of neurological or psychiatric disease based on the criteria of Diagnostic and Statistical Manual of Mental Disorders, Fourth Edition (DSM-IV) evaluated by experienced psychiatric physicians; 2) had a history of loss of consciousness for more than 5 min; 3) pregnant or lactating women or women planning to become pregnant; and 4) had contraindications to MRI scanning. All patients were diagnosed using the criteria from the structured clinical interview SCID-I/P (Structured Clinical Interview for DSM-IV Axis I Disorders, Patient Edition) manual, which was developed according to DSM-IV. Two psychiatrists confirmed that participants were in line with clinical diagnostic criteria of MDD, BD and SZ. All psychiatric patients were further excluded if they: 1) had suffered from other diagnoses of DSM-IV axis I and axis II; 2) had mental disorders (alcohol, drugs, etc.) caused by substance abuse, severe and unstable physical diseases, diagnosed diabetes, thyroid disease, hypertension, heart disease, etc.; 3) had narrow-angle glaucoma; 4) had a history of epilepsy, and those with high febrile convulsions; 5) had used non-convulsive electroconvulsive therapy 6 months before enrollment; 6) pregnant or lactating women or women planning to become pregnant; and 7) had contraindications to MRI scanning. The final analyzed sample included 409 healthy controls from the Healthy dataset, and 328 HC, 56 with MDD, 31 with BD, and 73 with schizophrenia from the Clinical dataset. The study has been approved by the ethics committees of Peking University Sixth Hospital and written informed consent was received from each subject prior to the formal testing.

### Image acquisition and preprocessing

Images data of all participants were acquired on a 3.0 T GE Discovery MR750 scanner in the Center for MRI Research, Peking University (Healthy dataset), and the Center for Neuroimaging, Peking University Sixth Hospital (Clinical dataset). The high-resolution structural T1-weighted MRI was acquired in a sagittal orientation using an axial 3D fast, spoiled gradient recalled (FSPGR) sequence with the following parameters: time repetition (TR) = 6.66 ms, time echo (TE) = 2.93 ms, field of view (FOV) = 256 × 256 mm^2^, slice thickness/gap = 1.0/0 mm, acquisition voxel size = 1 × 1 × 1mm^3^, flip angle = 12°, 192 contiguous sagittal slices. For the fMRI, each echo-planar image consisted of 33 (4.2 mm thick, 0 mm gap) axial slices covering the entire cerebrum and cerebellum (TR/TE = 2000/30 ms, flip angle = 90°, FOV = 224 × 224 mm^2^, 64 × 64 matrix). Scanning parameters were selected to optimize the stability and quality of the blood oxygenation level-dependent (BOLD) signal.

The preprocessing of functional imaging was preprocessed using the SPM12 (Statistical Parametric Mapping, <http://www.fil.ion.ucl.ac.uk/spm>) with a standardized protocol whose details can be found elsewhere (1). Specifically, image processing included 1) excluding the first 4 images as dummy scans; 2) performing slice-timing correction in remaining images; 3) realigning remaining images to the first volume for head motion correction using a six-parameter (rigid body) linear transformation; 4) conducting brain tissue segmentation and spatial normalization using the standard stereotaxic space of Montreal Neurological Institute (MNI) template; 5) performing the spatial smoothing with a Gaussian filter set at 8mm full-width at half-maximum. Each task-evoked stimulus was modeled as a separate delta function and convolved with a canonical hemodynamic response function, ratio normalized to the whole-brain global mean to control for systematic differences in global activity, and temporally filtered using a high-pass filter of 128s. Each task-evoked stimulus event was modeled for correctly performed trials. Incorrect responses and residual movement parameters were also modeled as regressors of no interest.

### Task paradigm of emotional episodic memory

Disruptions in emotional episodic memory has been proposed as a prominent type of affective symptoms and may be driven by various emotion-related neural dysfunctions (2). All participants in the Healthy dataset and the Clinical dataset performed a fMRI paradigm of emotional episodic memory task. The paradigm consisted of encoding and retrieval phases and each phase included neutral and aversive images selected from the International Affective Picture System (3, 4). Both encoding and retrieval phase comprised 4 blocks of neutral scenes and 4 blocks of aversive scenes which were continuously presented, and each block included 6 scenes for 3s each scene, alternating with 9 blocks of rest (18s of fixation). Within the scanner, participants firstly completed an encoding phase in which they were instructed to determine whether each image represented an ‘indoor’ or ‘outdoor’ scene and respond via a button press. The presentation order of ‘indoor’ and ‘outdoor’ scenes was randomized, and participants were not informed about the subsequent retrieval phase before scanning and thus were not realized that they were engaged in a memory task. About 2 minutes after the encoding phase, participants in retrieval phase need to memorize whether the image presented was seen during the encoding session and respond ‘new’ or for ‘old’ via a button press. During retrieval phase, half of the scenes were old (i.e., presented during encoding phase), and half were new (i.e., not presented during encoding phase). The presentation order of ‘new’ and ‘old’ scenes was also randomized. The order of the presentation of blocks (neutral and aversive scenes) in encoding and retrieval phase was counterbalanced across individuals. Finally, both reaction time (RT) and accuracy (ACC) in encoding and retrieval phase were extracted from log files of task performance after scanning, and response ACC was also estimated by sensitivity (d prime, d′) based on signal detection measures. The statistical analyses of task performances within group were used to paired-sample *t*-test to examine the difference between neutral and aversive conditions to investigate the memory-enhancing effect induced by emotional stimuli in each group. The one-way ANOVA was used to evaluate the between-group differences for emotional enhancing effect in RT and ACC of task performances.

Task performances of the Healthy and the Clinical datasets were summarized in Table S2. The statistical results showed that the memory-enhancing effect in the negative emotional condition was also observed in each group (HC_test_: *t* = 8.722; MDD: *t* = 5.094; BD: *t* = 4.543; SZ: *t* = 4.249; all *p* < .001). In addition, d′ values were statistically significantly greater during the retrieval phase compared to the encoding phase across all groups (HC_train_: *t* = 14.168; HC_test_: *t* = 10.658; MDD: *t* = 6.618; BD: *t* = 4.071; SZ: *t* = 5.184; all *p* < .001). There were significant between-group differences in encoding neutral RT (*F* = 16.724, *p* < .001) and aversive RT (*F* = 23.949, *p* < .001), as well as retrieval neutral RT (*F* = 9.928, *p* < .001) and aversive RT (*F* = 14.672, *p* < .001) across HC, MDD, BD and SZ groups. In contrast, no significant differences were observed in retrieval neutral ACC (*H* = 4.751, *p* = .191) and aversive ACC (*H* = 5.028 *p* = .166) among the HC and all case groups. There is also no significant difference was found in retrieval ACC change (aversive − neutral) across HC, MDD, BD and SZ groups (*F* = 0.117, *p* = .950).

### Preprocessing of brain-wide gene expression dataset

The brain-wide gene expression data were sourced from six neurotypical adult donors provided by the Allen Human Brain Atlas (AHBA, <http://human.brain-map.org>). The microarray data derived from post-modern brain tissue samples underwent preprocessing via the abagen toolbox (<https://www.github.com/netneurolab/abagen>), following established protocols (5). Specifically, genetic probes were reannotated based on guidelines from previously published studies (5). Probes with intensity values below the background noise threshold, set at 50%, were excluded through intensity-based filtering. Spatial registration of each tissue sample to Montreal Neurological Institute (MNI) coordinate space was performed using T1-weighted images from each donor. Brain regions were assigned to tissue samples based on their corresponding MNI coordinates (<https://github.com/chrisfilo/alleninf>). Due to the limited availability of right hemisphere data (only two donors), the analysis of gene expression was restricted to 41 left hemisphere regions. Only one brain region lacking corresponding tissue samples was interpolated. For each donor, gene expression values across brain regions were normalized using a robust sigmoid function and subsequently rescaled to the unit interval. This preprocessing resulted in a gene expression map (41 regions × 15,633 genes), which was utilized for subsequent imaging-transcriptome association analyses.

### Statistical analysis for group convergent effect

The statistical analyses for group convergent effect of individual deviations derived from normative modelling were conducted for HC_test_ and each case in the Clinical dataset across the regional, network and circuit level, aiming at group differences after removing intersubject heterogeneity go beyond the evaluation of traditional group-averaged differences. The pipeline of detailed statistical analysis was shown as Figure S2.

The group convergent effect in the regional level were conducted for HC_test_ and each case in the Clinical dataset and assessed by using a nonparametric group-based permutation test. Specifically, the percentage maps showing extreme individual deviation in HC_test_­ were subtracted from each case’s percentage map, resulting in percentage difference maps (Δ percentage map) of each disorder at any given brain region. We permuted group labels of case group and HC group, and repeated the procedure 10,000 times to re-calculated the group-specific percentage map and percentage difference (Δ percentage map). We then derived an empirical distribution of percentage difference maps under the null hypothesis of random group assignment. The *p* values were obtained as the proportion of null values that exceeded the observed difference and statistical significance was set at 0.05 (over 95% proportion) and correction for multiple comparisons was performed using the false discovery rate (*p* < .05, FDR corrected).

To characterize emotional effect at the network level, we positioned four brain emotional networks (ENs) (6) on DK atlas (Figure S1). These four ENs included a total of 36 brain areas derived from DK atlas and exhibited distinct spatial distributions and functional profiles were included in these networks (Table S1). At network level, we characterized network deviations by assigning each region to one of four ENs. The network percentage showing an extreme deviation was estimated as the proportion of individuals that showed at least one extreme deviation in a region within the network, separately for positive and negative extrema. The case-specific network difference then was evaluated by computing differences between network percentage of HC_test_­ and each case. The statistically significance of network difference (Δ network percentage map) was assessed by 10,000 times group-based permutation test, as at regional level. The statistically significance defined using a threshold at 0.05 and adjusted by FDR method (*p* < .05, FDR corrected).

The network-level analysis preliminary explored individualized representation of activated deviations at the level of more extensive emotional networks. We further investigated whether the convergence of regional deviation in emotional networks could be driven by functional segregation and integration within or between networks. Thus, we estimated circuit level individual deviations using lesion network mapping methodology (7, 8). Firstly, each region showing an extreme deviation was mapped into functional circuitry defined by psychophysiological interaction (PPI) analysis in each group. The psychophysiological interaction (PPI) was calculated in any individual of HC_test_ and each case group to define the functional circuitry of each region showing an extreme deviation under the “aversive − neutral” condition (see “psychophysiological interaction under “aversive − neutral” condition”). Specifically, each deviant region was used as a seed and contrast its task-based functional connectivity maps with other brain regions, which was determined by PPI analysis in HC and each case group. Then, we thresholded the individualized deviation of functional circuitry maps according to statistical significance across all participants of each group using one-way *t*-test (*p* < .0167 (.05/3), account for 3 group analyses). By binarizing the value of statistically significant functional circuitry as 1, extreme circuit deviations of each participant for each group were obtained at any regions survived the statistical threshold and took the union of the extreme circuit maps across that set of seeds in each person. The union of the extreme functional circuit in each person were obtained for positive and negative deviations, respectively, as final extreme circuit deviation. Finally, a group-specific circuit percentage map was quantified as the proportion of individuals showing an extreme circuit deviation was calculated in positive and negative aspects. Following the analysis of regional level, the circuit percentage map was yielded by calculating the proportion of individuals in each extreme circuit deviation, and the circuit percentage maps in HC_test_­ were subtracted from each case’s percentage map and contributed to circuit percentage difference maps (Δ circuit percentage map) of clinical group relative to the HC_test_ group. The significance of circuit percentage difference maps was evaluated by permuting the group labels of the individual-specific circuit percentage maps 10,000 times, using a threshold of *P* < .05 with FDR correction. The significant seed-based functional circuit was further located into four ENs to assess the mediate role of circuit convergent effect in relationship between individual deviations at regional level and network level.

### The model generalizability of normative modelling

The internal evaluation of model generalizability of normative functioning underlying emotional episodic memory was assessed using 10-fold cross-validation applied to HC_train_, and replicated estimation in HC_test_­. We assessed performance for the normative models for each brain region by evaluating the standardized mean-squared error (SMSE) and the mean standardized log-loss (MSLL) in HC_train_ and HC_test_ cohort, respectively. As external validation, we positioned each participant in HC_test_ within the normative range and further calculated the subject-level distribution similarity between HC_train_ and HC_test_ cohort using Kolmogorov-Smirnov test and the spatial-level distribution similarity using spatial correction analysis with spatial autocorrelation corrected. Statistic threshold was set at 0.05. As shown in Figure S3, a high internal generalizability of normative modelling was exhibited, showing with a SMSE close to 1 (HC_train_: 1.050 ± 0.030; HC_train_: 1.047 ± 0.031) and a MSLL close to 0 (HC_train_: 0.029 ± 0.016; HC_train_: 0.023 ± 0.026). For external validation, as shown in Figure 2 in main text, there was no significant difference of the distribution of mean regional normative *z* scores between HC_train_ and HC_test_ (Kolmogorov-Smirnov test, *K-S* = 0.087, *p* = .320). This spatial distribution of emotional memory-related brain activation is highly correlated with individual-level mean distribution patterns of HC_train_ (*r* = 0.973, *p*_spin_ < .001, 10,000 times permutation tests with spatial autocorrelation), and mean regional deviation map in HC_train_ and HC_test_ was highly associated (Figure 2C, *r* = 0.956, *p*_spin_ < .001, 10,000 times permutation tests with spatial autocorrelation).

### Overview of measurement stability of normative modelling

To demonstrate the stability of individualized measurement defined from normative modelling, we validated our results by considering several potential confounding factors. Firstly, we re-constructed the normative models by including sex and age into predictor set to evaluate potential age/sex effects on model design. Secondly, we evaluated the influence of different brain parcellation atlas, including Anatomical Automatic Labelling (<https://www.gin.cnrs.fr/en/tools/aal/>) and Brainnetome (<http://atlas.brainnetome.org/>). Thirdly, to validate whether our results could be influenced by the medication effect, the correlation analysis between individualized regional deviations and dose of medicine was performed at each disorder. The results of different analyses of measurement stability were shown in the following section.

### Measurement stability analysis 1: age and sex effects on normative model

Age and age are widely used clinically covariables to construct normative models, to further depict the age trend under different sex conditions in prior studies. To evaluate potential age and sex effects on model design, we also constructed the normative models by including sex and age into original predictor set (behaviour performance comprising of reaction times in encoding and retrieval phase and accuracy on retrieval phase) in HC_train_ and HC_test._ The spatial correlation was conducted between spatial distributions including age and sex and not including them. The statistically significance was defined using a threshold of *p*_spin_ < .05 with 10,000 times spherical rotations-based spatial autocorrelation. The results showed that there are significant spatial correlations between four different conditions (all *p*_spin_ < .001, Figure S4), implying that sex and age had no significant effect on our normative model.

### Measurement stability analysis 2: different brain parcellation atlas

The percentage differences at each disorder were re-evaluated using different brain parcellations, including Anatomical Automatic Labelling (AAL) atlas and Brainnetome atlas. We applied correlation analysis to assess the correlation between mean individualized deviations was used to examine the repeatability across atlas. The results showed that the spatial distribution pattern of percentage differences in each atlas exhibited high similarity and mean deviations in AAL and Brainnetome atlas were both statistically significantly correlated with our main findings using DK atlas in MDD (AAL: *r* = 0.997, *p* < .001; Brainnetome: *r* = 0.995, *p* < .001; Figure S5A), BD (AAL: *r* = 0.994, *p* < .001; Brainnetome: *r* = 0.995, *p* < .001; Figure S5B), and SZ patients (AAL: *r* = 0.998, *p* < .001; Brainnetome: *r* = 0.996, *p* < .001; Figure S5C).

### Measurement stability analysis 3: the influences of medication effect

In the Clinical dataset of current study, the 66.07% (37/56) of patients with MDD and 53.42% (39/73) of patients with SZ are taking medicine. The dose of medicine in MDD and SZ patients was transformed to Fluoxetine equivalent and Olanzapine equivalent, respectively. In BD patients, 44.16% (14/31) of patients with BD are taking Sodium valproate, 29.03% (9/31) Lithium carbonate and 22.58% (7/31) Quetiapine. To evaluate the influences of medication effect, the Pearson correlation analysis was conducted between dose of medicine and individualized regional deviations in the patients with MDD, BD and SZ. The statistically significance was defined using a threshold of *p* < .05 and adjusted by FDR. As shown in Figure S6, there are no significant correlation between dose of medicine and individual regional deviations in MDD (*p* < .05; FDR corrected; Figure S6A), BD (*p* < .05; FDR corrected; Figure S6B) and SZ patients (*p* < .05; FDR corrected; Figure S6C).

### Clinical heterogeneity of functional activations deviations in each disorder

Regions with extreme deviations exhibited considerable intersubject heterogeneity in patients with psychiatric disorders. There were 44.64% of the patients with MDD showing infra-normal or supra-normal regions in at least one brain region, including infra-normal regions in 25.00% of patients and supra-normal regions in 21.43% of patients. Regionally, a total of 92.68% of brain regions showed infra-normal or supra-normal in at least one patient, with infra-normal regions in 40.24% and supra-normal regions in 90.24%. However, for any single brain region, the percentage of individuals with infra-normal regions was less than 10.71% and with supra-normal regions was less than 8.93% (Figure S7A&D). More than 64.52% of BD patients were exhibited infra-normal or supra-normal regions in at least one brain region (infra-normal regions: 54.84%; supra-normal regions: 12.90%). From the perspective of brain regions, 80.49% of brain regions showed infra-normal or supra-normal in at least one patient, showing with 65.85% in infra-normal and 37.80% in supra-normal patients. In contrast, there are less than 25.81% and 9.68% of percentage of individuals with infra-normal and supra-normal at single region, respectively (Figure S7B&E). For SZ patients, the percentage of individuals with extreme deviations in at least one brain region was 54.79% (infra-normal regions: 28.77%; supra-normal regions: 30.14%). A total of 100% of brain regions showed extreme deviations in at least one patient, with 89.02% in infra-normal and 93.90% in supra-normal. Nevertheless, the percentage of individuals at specific brain region level was less than 8.22% in infra-normal regions and less than 6.85% in supra-normal regions (Figure S7C&F).

### Traditional group-level comparison using general linear model

We examine case-control difference of functional activation under “aversive > neutral” condition using traditional group-level comparison based on general linear model. This model was conducted in the Clinical dataset to explore the differences of functional activation of MDD, BD and SZ patients relative to HC_test_. For each comparison, the effect of sex and age were set as covariables to control. Statistical significance was set at 0.05 and correction for multiple comparisons was performed using the false discovery rate (*p* < .05, FDR corrected). No significant differences were found in MDD (Figure S6A), BD (Figure S6B) and SZ patients (Figure S6C) compared with HC_test_.

### Shared regional differences across MDD, BD and SZ patients

The intersection of regions with infra-normal or supra-normal across different disorders was obtained to represent shared regional differences in emotional episodic memory. The shared regions across MDD, BD and SZ patients included left insula, right supramarginal gyrus, and right amygdala, which characterized a transdiagnostic functional network of negativity bias in emotional memory. Additionally, the shared regions across MDD and BD patients were consisted of right pars orbitalis and parahippocampal gyrus, and the shared regions across MDD and SZ patients were in left bankssts, left lingual gyrus, right pars opercularis, and right superior frontal gyrus. The shared regions across BD and SZ patients exhibited more regions with the same alterations, mainly including insular cortex, temporal gyrus, striatum, and hippocampus (Figure S7). The shared regions were summarized in Table S3.

### Threshold-weighted deviation analysis

Following a pervious study, we used a threshold-weighted deviation analysis to estimate the effect on the results of threshold for defining extreme deviations (9). Specifically, we firstly set the threshold range of from 1.64 to 3.10 with 100 equal log-space increments, separately for positive and negative deviation. Then, for each group, we respectively calculated the individual extreme deviation and obtained a region- and threshold-specific percentage map (the number of regions × the number of threshold) quantifying the proportion of participants with extreme deviation in each case and control group. Next, a threshold-weighted function according to a previous study was applied to penalize fewer conservative thresholds, which ensured that values of percentage map range between 0 and 1 and as followed:

$$W_{threshold}=2 \times\frac{threshold-min(threshold)}{\max\left( threshold \right)-min(threshold)}$$

Finally, the region- and threshold-specific percentage map was transformed to a threshold-weighted percentage map by taking the area under the curve of the cumulative histogram of the proportion of individuals at each threshold for each region.

The spatial correlation was conducted between percentage difference maps thresholded at 2.6 and thresholded using the above-mentioned threshold-weighted deviation analysis at all case groups. The statistically significance was defined using a threshold of *p*_spin_ < .05 with 10,000 times spherical rotations-based spatial autocorrelation. As shown in Figure S10, we found that there are significant spatial correlations between different threshold setting in case-control percentage differences of MDD (infra-normal: *r* = 0.920, *p*_spin_ < .001; supra-normal: *r* = 0.918, *p*_spin_ < .001; Figure S10A), BD (infra-normal: *r* = 0.961, *p*_spin_ < .001; supra-normal: *r* = 0.957, *p*_spin_ < .001; Figure S10B) and SZ (infra-normal: *r* = 0.927, *p*_spin_ < .001; supra-normal: *r* = 0.930, *p*_spin_ < .001; Figure S10C).

### Large-scale emotional networks and its association with functional activation

Extreme deviations of brain regions were also embedded into large-scale emotional network with specific emotional profiling to explore the relevant emotional functional disruptions in distinct psychiatric disorders. A well-recognized framework of four large-scale emotional networks (ENs) with distinct spatial distributions and functional profiles was revealed by a recent meta-analysis study (6), and positioned on DK atlas (Figure S1). A total of 36 brain areas were included in these four ENs in current study (Table S2). To examine the significant functional activation at “aversive > neutral” condition, we conducted one-way *t*-tests at healthy controls from the Healthy dataset (HC_train_). Statistical significance was set at 0.05 and correction for multiple comparisons was performed using the false discovery rate (*p* < .05, FDR corrected). Furthermore, we computed the percentage of regions showing statistically significant activation in different ENs to evaluate the rationality of using these four emotional networks.

The results showed that “aversive > neutral” condition of emotional memory task exhibited a broad significant functional activation in HC_train_ group (Figure S11). The regions with statistically significantly positive activation were mainly located in with the occipital and temporal cortex, while those regions with significant negative activation were mostly in frontal cortex, insula and subcortex (Figure S11A). The percentage of ENs-related regions showing signification activation is high at total ENs (percentage of total ENs = 90.32%) and different subnetwork of ENs (percentage of total EN 1 = 90.32%; percentage of total EN 2 = 100%; percentage of total EN 3 = 100%; percentage of total EN 4 = 77.78%) (Figure S11B). These results suggested that regions with functional activation at emotional memory task may be embedded at the level of the broader emotional networks.

### Feature weight analysis of predicted model in affective symptom association

Following the machine-learning model that investigate the relationships between individualized neurofunctional deviations and clinical affective symptoms in different diagnostic groups, feature weight of each region on predicted model was estimated using virtual lesion analysis. This method based on a re-calculated performance (correlation coefficient between observed scores and predicted scores) when removed from the predictive model at each iteration (i.e., virtual lesion) was used to determine the important regions and networks, respectively. If the prediction accuracy of model decreased after removing a region, we considered the region to be important for prediction. As shown in Figure S12, the important features have been identified in those emotional networks specific to heterogenous deviations at each psychiatric disorder: EN 3 for MDD, EN 4 for BD, and EN 1 for SZ (Figure S12).

### Psychophysiological interaction under “aversive > neutral” condition

The psychophysiological interaction (PPI) analysis under “aversive > neutral” condition was used to evaluate the functional circuitry in any individual of HC_test_ and each case group. Following previous studies (10, 11), the BOLD signals for each participant under emotional episodic memory task were averaged in each brain regions defined by the Desikan-Killiany (DK) (12). Serial correlations then were removed using a first-order autoregressive model and a high-pass filter (128 s) was applied to remove low-frequency noise (13, 14). We performed a general linear model (GLM) to predict functional activation of brain regions other than seed regions as a function of the activation of seed regions (the “physiological” regressor), effect of task under “aversive > neutral” condition convolved with the hemodynamic response function (the “psychological” regressor) and the PPIs of these tasks with the deconvolved signal of the seed regions (the “PPI” regressor). The β coefficient associated with PPI under “aversive > neutral” condition correspond to the modulation of functional connectivity for task in HC_test_ and each disorder.

### Group convergence effect of individual deviation at circuit level

Our analysis showed that regional deviations underlying affective symptoms of distinct disorders couple into dissociable emotional networks. However, network analysis only extends our understanding to arbitrarily grouped regions and may not reflect inter- and intra-network functional interaction. To address this, we conducted a circuit-level analysis to estimate whether regional activation deviations and broader deviant patterns at the network level were mediated by functional couplings with deviant regions. Compared to HC, MDD patients showed infra-normal seed-based functional circuits predominately in the prefrontal cortex and visual regions and supra-normal functional circuit in the fusiform gyrus and medial orbitofrontal gyrus (*p* < .0167 (.05/3), FDR corrected; Figure S14A; Table S8). The most significant circuit were located in EN 3 (*p* < .000203 (.05/3/82), Bonferroni corrected; Figure S14B). In BD patients, infra-normal circuits were mainly in the occipital cortex and supramarginal gyrus, while supra-normal circuits were in the cingulate gyrus, thalamus, and striatum (*p* < .0167 (.05/3), FDR corrected; Figure S14C; Table S9), with the most infra-normal regions in EN 4 (Bonferroni corrected; Figure S14D). SZ patients exhibited significant infra-normal circuit in the parietal and frontal gyri, and large-scale supra-normal circuits involving frontal, temporal and occipital regions (*p* < .0167 (.05/3), FDR corrected; Figure S14E; Table S10). The most significant infra-normal regions were in EN 1 (Figure S14F). Collectively, these findings align with the network-level analysis results. Our findings further revealed that the convergence of functional deviations at the circuit-level suggested that dysfunctional interactions within specific emotional networks may drive disrupted patterns underlying affective symptoms in distinct mental illnesses. In patients with MDD, excessive information integration within the network of emotion perception and generation mediated its functional hyper-activation. Medial orbitofrontal gyrus is the main disrupted hub, which is considered to participate in the generation of negative emotions and the monitoring of depression in the past research (15), and meanwhile play the role of emotional regulation (16, 17). Patients with BD is more related to the dysfunction of emotional network responsible for internal feelings and emotional states, including IPG, precuneus and middle occipital gyrus. Among them, the middle occipital gyrus was identified as the impaired core of excessive functional segregation within network. In a recent study, this region was thought to be related to the semantic and conceptual representations of negative stimuli (18), and its dysfunction may explain emotional state caused by the inability to recognize emotional meaning. Consistent with the previous research hypothesis, our findings also believed that the affective symptoms of SZ stems from the impaired cognitive controls (19). The decrease of functional integration in dorsal emotional regulation system with lateral prefrontal cortex and inferior parietal gyrus as the core may partly explained the defect of emotional cognitive control in SZ, which leads to the weakening of the inhibition of amygdala functional activities.

### Supplementary Tables

#### Table S1. Large-scale emotional networks

|  |  |  | MNI peak coordinates | | |
| --- | --- | --- | --- | --- | --- |
| Networks | Region name | DK | x | y | z |
| EN 1 | Superior frontal gyrus (L) | 27 | 0 | 24 | 50 |
|  | Caudal middle frontal gyrus (R) | 44 | 40 | 24 | 42 |
|  | Inferior parietal gyrus (R) | 48 | 58 | − 52 | 38 |
|  | Inferior parietal gyrus (L) | 7 | − 58 | − 50 | 44 |
|  | Rostral middle frontal gyrus (L) | 26 | − 36 | 52 | − 2 |
|  | Caudal middle frontal gyrus (L) | 3 | − 42 | 14 | 48 |
|  | Rostral middle frontal gyrus (R) | 67 | 42 | 46 | − 8 |
|  | Insula (R) | 75 | 36 | 16 | 6 |
|  | Posterior cingulate (R) | 63 | 2 | − 22 | 30 |
|  | Precuneus (R) | 65 | 10 | − 64 | 36 |
| EN 2 | Pars orbitalis (L) | 18 | − 46 | 24 | − 8 |
|  | Superior frontal gyrus (L) | 27 | − 4 | 10 | 62 |
|  | Pars triangularis (R) | 60 | 50 | 28 | − 8 |
|  | Superior temporal gyrus (L) | 29 | − 54 | − 34 | − 2 |
|  | Middle temporal gyrus (L) | 14 | − 44 | 6 | 50 |
|  | Caudal middle frontal gyrus (L) | 3 | − 30 | 48 | 26 |
|  | Rostral middle frontal gyrus (L) | 26 | − 16 | 10 | 12 |
|  | Caudate (L) | 36 | 36 | − 60 | − 30 |
| EN 3 | Amygdala (L) | 40 | − 22 | − 4 | − 16 |
|  | Amygdala (R) | 81 | 24 | − 4 | − 18 |
|  | Fusiform gyrus (R) | 47 | 40 | − 46 | − 18 |
|  | Thalamus (R) | 76 | 6 | − 26 | 0 |
|  | Fusiform gyrus (L) | 6 | − 38 | − 54 | − 14 |
|  | Parahippocampal gyrus (L) | 15 | − 22 | − 28 | − 4 |
|  | Medial orbitofrontal gyrus (L) | 13 | 0 | 54 | −10 |
|  | Medial orbitofrontal gyrus (R) | 54 | 0 | 54 | −10 |
|  | Lingual gyrus (L) | 12 | − 42 | − 76 | − 6 |
| EN 4 | Postcentral gyrus (L) | 21 | − 58 | − 22 | 32 |
|  | Insula (L) | 34 | − 44 | − 4 | 10 |
|  | Superior parietal gyrus (L) | 28 | − 28 | − 52 | 56 |
|  | Postcentral gyrus (R) | 62 | 62 | − 22 | 30 |
|  | Cuneus (L) | 4 | − 10 | − 76 | 22 |
|  | Lateral occipital gyrus (L) | 10 | − 48 | − 74 | 2 |
|  | Thalamus (R) | 76 | 10 | − 26 | − 4 |
|  | Superior parietal gyrus (R) | 69 | 28 | − 60 | 38 |
|  | Precuneus (R) | 65 | 16 | − 56 | 16 |

**Abbreviations**: DK, Desikan-Killiany atlas; L, left; R, right

#### Table S2. The task performances of healthy and clinical datasets

|  | Healthy dataset |  | Clinical dataset | | | | |
| --- | --- | --- | --- | --- | --- | --- | --- |
|  | HC_trian_  (n = 409) |  | HC_test_  (n = 164) | MDD  (n = 56) | BD  (n = 31) | SZ  (n = 73) | Statistical test  *F* / *H* (*p*) |
| Encoding neutral RT, ms | | | | | | | 16.724 (< .001^a^) |
|  | 833.069 (143.190) |  | 820.779 (133.333) | 850.007 (157.554) | 886.614 (184.863) | 980.024 (209.555) | HC = MDD = BD < SZ^c^ |
| Encoding aversive RT, ms | | | | | | | 23.949 (< .001^a^) |
|  | 1012.998 (182.446) |  | 1020.317 (180.630) | 1121.826 (229.932) | 1196.299 (358.112) | 1282.910 (239.837) | HC < MDD = BD < SZ^c^ |
| Retrieval neutral RT, ms | | | | | | | 9.928 (< .001^a^) |
|  | 1067.747 (155.136) |  | 1025.032 (159.672) | 1051.631 (140.938) | 1107.519 (165.464) | 1152.416 (217.534) | HC = MDD = BD < SZ^c^ |
| Retrieval aversive RT, ms | | | | | | | 14.672 (< .001^a^) |
|  | 1098.223 (163.875) |  | 1061.150 (153.655) | 1093.458 (184.097) | 1221.292 (187.848) | 1196.758 (199.159) | HC = MDD < BD = SZ^c^ |
| Retrieval neutral ACC | | | | | | | 4.751  (.191^b^) |
|  | 0.856  (0.079) |  | 0.849 (0.084) | 0.839 (0.090) | 0.835 (0.071) | 0.816 (0.112) |  |
| Retrieval aversive ACC | | | | | | | 5.082  (.166^b^) |
|  | 0.918  (0.065) |  | 0.920 (0.062) | 0.918 (0.074) | 0.915 (0.068) | 0.891 (0.099) |  |
| Retrieval ACC change (aversive − neutral) | | | | | | | 0.117  (.950^c^) |
|  | 0.062  (0.089) |  | 0.071 (0.090) | 0.079 (0.100) | 0.080 (0.108) | 0.075 (0.137) |  |

Data are presented as mean (standard deviations). HC, healthy control; MDD, major depressive disorder; BD, bipolar disorder; SZ, schizophrenia; RT, reaction time; ACC, accuracy

^a^ The *p*-value was obtained by a one-way ANOVA with post hoc analyses.

^b^ The *p*-value was obtained by a Kruskal-Wallis test.

^d^ Post hoc comparison.

#### Table S3. Shared regional differences across all cases

| Merged cases | Direction | Direction |
| --- | --- | --- |
| MDD ∩ BD | Infra-normal |  |
|  | Supra-normal | Insula (L) |
|  |  | Parahippocampal gyrus (R) |
|  |  | Pars orbitalis (R) |
|  |  | Supramarginal gyrus (R) |
|  |  | Amygdala (R) |
| MDD ∩ SZ | Infra-normal |  |
|  | Supra-normal | Bankssts (L) |
|  |  | Lingual gyrus (L) |
|  |  | Pars opercularis (L) |
|  |  | Insula (L) |
|  |  | Superior frontal gyrus (R) |
|  |  | Supramarginal gyrus (R) |
|  |  | Amygdala (R) |
| BD ∩ SZ | Infra-normal | Caudate (L) |
|  |  | Hippocampus (L) |
|  |  | Insula (R) |
|  |  | Caudate (R) |
|  |  | Amygdala (R) |
|  | Supra-normal | Transverse temporal gyrus (L) |
|  |  | Insula (L) |
|  |  | Putamen (L) |
|  |  | Supramarginal gyrus (R) |
|  |  | Insula (R) |
|  |  | Amygdala (R) |
| MDD ∩ BD ∩ SZ | Infra-normal |  |
|  | Supra-normal | Insula (L) |
|  |  | Supramarginal gyrus (R) |
|  |  | Amygdala (R) |

MDD, major depressive disorder; BD, bipolar disorder; SZ, schizophrenia; L, left; R, right

#### Table S4. Regional differences between MDD and HC_test_

| Direction | Regions | Δ percentage | *p* |
| --- | --- | --- | --- |
| Infra-normal |  |  |  |
|  | Entorhinal gyrus (L) | 4.75% | .007 |
|  | Fusiform gyrus (L) | − 3.05% | < .001 |
|  | Lateral orbitofrontal gyrus (L) | − 3.66% | < .001 |
|  | Precentral gyrus (L) | − 2.44% | < .001 |
|  | Rostral anterior cingulate (L) | − 2.44% | < .001 |
|  | Superior frontal gyrus (L) | − 3.05% | < .001 |
|  | Transverse temporal gyrus (L) | − 2.44% | < .001 |
|  | Pallidum (L) | − 1.83% | < .001 |
|  | Accumbens (L) | − 1.83% | < .001 |
|  | Bankssts (R) | − 2.44% | < .001 |
|  | Fusiform gyrus (R) | − 1.83% | < .001 |
|  | Lateral orbitofrontal gyrus (R) | − 2.44% | < .001 |
|  | Rostral anterior cingulate (R) | − 1.83% | < .001 |
|  | Superior temporal gyrus (R) | 1.79% | < .001 |
| Supra-normal |  |  |  |
|  | Bankssts (L) | 3.57% | < .001 |
|  | Lingual gyrus (L) | 5.36% | < .001 |
|  | Pars opercularis (L) | 8.32% | < .001 |
|  | Posterior cingulate (L) | 3.57% | < .001 |
|  | Rostral anterior cingulate (L) | 4.75% | .016 |
|  | Superior frontal gyrus (L) | 5.36% | < .001 |
|  | Superior temporal gyrus (L) | 4.75% | .013 |
|  | Supramarginal gyrus (L) | 5.36% | < .001 |
|  | Insula (L) | 3.57% | < .001 |
|  | Parahippocampal gyrus (R) | − 2.44% | < .001 |
|  | Pars orbitalis (R) | 4.75% | .016 |
|  | Posterior cingulate (R) | 3.57% | < .001 |
|  | Superior frontal gyrus (R) | 4.75% | .016 |
|  | Superior temporal gyrus (R) | 4.75% | .016 |
|  | Supramarginal gyrus (R) | 3.57% | < .001 |
|  | Amygdala (R) | 5.36% | < .001 |

Δ percentage is obtained by subtracting percentage maps of HC_test_­ from percentage map of MDD patients. The threshold of statistical significance was set at 0.05 and correction for multiple comparisons was performed using the false discovery rate (*p* < 0.05/3, FDR corrected). HC, healthy control; MDD, major depressive disorder; L, left; R, right

#### Table S5. Regional differences between BD and HC_test_

| Direction | Regions | Δ percentage | *p* |
| --- | --- | --- | --- |
| Infra-normal |  |  |  |
|  | Lateral occipital gyrus (L) | 19.53% | < .001 |
|  | Precuneus (L) | 5.84% | .011 |
|  | Frontal pole (L) | 11.86% | .014 |
|  | Caudate (L) | 3.23% | < .001 |
|  | Hippocampus (L) | 5.84% | .008 |
|  | Bankssts (R) | 13.69% | .002 |
|  | Inferior temporal gyrus (R) | 22.76% | < .001 |
|  | Lateral occipital gyrus (R) | 11.07% | .003 |
|  | Precuneus (R) | 6.45% | < .001 |
|  | Superior parietal gyrus (R) | 8.46% | .005 |
|  | Insula (R) | 3.23% | < .001 |
|  | Caudate (R) | 3.23% | < .001 |
|  | Amygdala (R) | 6.45% | < .001 |
| Supra-normal |  |  |  |
|  | Transverse temporal gyrus (L) | 3.23% | < .001 |
|  | Insula (L) | 9.68% | < .001 |
|  | Putamen (L) | 5.84% | .005 |
|  | Caudal anterior cingulate (R) | − 3.05% | < .001 |
|  | Cuneus (R) | − 2.44% | < .001 |
|  | Parahippocampal gyrus (R) | 6.45% | < .001 |
|  | Pars orbitalis (R) | − 2.44% | < .001 |
|  | Supramarginal gyrus (R) | 5.84% | 0.005 |
|  | Insula (R) | 6.45% | < .001 |
|  | Caudate (R) | 5.84% | 0.005 |
|  | Amygdala (R) | 3.23% | < .001 |

Δ percentage is obtained by subtracting percentage maps of HC_test_­ from percentage map of BD patients. The threshold of statistical significance was set at 0.05 and correction for multiple comparisons was performed using the false discovery rate (*p* < 0.05/3, FDR corrected). HC, healthy control; BD, bipolar disorder; L, left; R, right

#### Table S6. Regional differences between SZ and HC_test_

| Direction | Regions | Δ percentage | *p* |
| --- | --- | --- | --- |
| Infra-normal |  |  |  |
|  | Inferior parietal gyrus (L) | 4.26% | 0.034 |
|  | Superior parietal gyrus (L) | − 2.44% | < .001 |
|  | Caudate (L) | 4.11% | < .001 |
|  | Hippocampus (L) | 6.24% | .004 |
|  | Parahippocampal gyrus (L) | 2.74% | < .001 |
|  | Insula (R) | 2.74% | < .001 |
|  | Caudate (R) | 5.48% | < .001 |
|  | Amygdala (R) | 4.11% | < .001 |
| Supra-normal |  |  |  |
|  | Bankssts (L) | 2.74% | < .001 |
|  | Fusiform gyrus (L) | 4.87% | .010 |
|  | Isthmus cingulate (L) | 4.87% | .011 |
|  | Lingual gyrus (L) | 4.11% | < .001 |
|  | Pars opercularis (L) | 4.87% | .011 |
|  | Transverse temporal gyrus (L) | 2.74% | < .001 |
|  | Insula (L) | 4.11% | < .001 |
|  | Putamen (L) | 4.87% | .010 |
|  | Accumbens (L) | 4.11% | < .001 |
|  | Superior frontal gyrus (R) | 2.74% | < .001 |
|  | Supramarginal gyrus (R) | 4.87% | .010 |
|  | Insula (R) | 4.11% | < .001 |
|  | Amygdala (R) | 4.11% | < .001 |
|  | Accumbens (R) | 5.48% | < .001 |

Δ percentage is obtained by subtracting percentage maps of HC_test_­ from percentage map of SZ patients. The threshold of statistical significance was set at 0.05 and correction for multiple comparisons was performed using the false discovery rate (*p* < 0.05/3, FDR corrected). HC, healthy control; SZ, schizophrenia; L, left; R, right

#### Table S7. Sample size evaluations

| Measures | Variable | Sample size |
| --- | --- | --- |
| Task performances | Encoding neutral RT | 124 |
|  | Encoding aversive RT | 128 |
|  | Retrieval neutral RT | 196 |
|  | Retrieval aversive RT | 112 |
|  | Retrieval neutral ACC | 804 |
|  | Retrieval aversive ACC | 776 |
| Functional neural activation | Bankssts (L) | 1832 |
|  | Caudal anterior cingulate gyrus (L) | 1108 |
|  | Caudal middle frontal gyrus (L) | 1628 |
|  | Cuneus (L) | 628 |
|  | Entorhinal gyrus (L) | 1868 |
|  | Fusiform gyrus (L) | 288 |
|  | Inferior parietal gyrus (L) | 2280 |
|  | Inferior temporal gyrus (L) | 492 |
|  | Isthmus cingulate gyrus (L) | 972 |
|  | Lateral occipital gyrus (L) | 148 |
|  | Lateral orbitofrontal gyrus (L) | 3620 |
|  | Lingual gyrus (L) | 744 |
|  | Medial orbitofrontal gyrus (L) | 1956 |
|  | Middle temporal gyrus (L) | 956 |
|  | Parahippocampal gyrus (L) | 8648 |
|  | Paracentral gyrus (L) | 660 |
|  | Pars opercularis (L) | 1688 |
|  | Pars orbitalis (L) | 1776 |
|  | Pars triangularis (L) | 1544 |
|  | Pericalcarine gyrus (L) | 968 |
|  | Postcentral gyrus (L) | 1224 |
|  | Posterior cingulate gyrus (L) | 4660 |
|  | Precentral gyrus (L) | 2452 |
|  | Precuneus | 860 |
|  | Rostral anterior cingulate gyrus (L) | 1820 |
|  | Rostral middle frontal gyrus (L) | 2608 |
|  | Superior frontal gyrus (L) | 1648 |
|  | Superior parietal gyrus (L) | 2288 |
|  | Superior temporal gyrus (L) | 2924 |
|  | Supramarginal gyrus (L) | 1684 |
|  | Frontal pole (L) | 872 |
|  | Temporal pole (L) | 1472 |
|  | Transverse temporal gyrus (L) | 824 |
|  | Insula (L) | 5520 |
|  | Thalamus (L) | 3856 |
|  | Caudate (L) | 7232 |
|  | Putamen (L) | 2372 |
|  | Pallidum (L) | 1344 |
|  | Hippocampus (L) | 1036 |
|  | Amygdala (L) | 1016 |
|  | Accumbens (L) | 396 |
|  | Bankssts (R) | 312 |
|  | Caudal anterior cingulate gyrus (R) | 1016 |
|  | Caudal middle frontal gyrus (R) | 1188 |
|  | Cuneus (R) | 612 |
|  | Entorhinal gyrus (R) | 2388 |
|  | Fusiform gyrus (R) | 336 |
|  | Inferior parietal gyrus (R) | 5640 |
|  | Inferior temporal gyrus (R) | 256 |
|  | Isthmus cingulate gyrus (R) | 888 |
|  | Lateral occipital gyrus (R) | 188 |
|  | Lateral orbitofrontal gyrus (R) | 1468 |
|  | Lingual gyrus (R) | 764 |
|  | Medial orbitofrontal gyrus (R) | 1756 |
|  | Middle temporal gyrus (R) | 1060 |
|  | Parahippocampal gyrus (R) | 1880 |
|  | Paracentral gyrus (R) | 828 |
|  | Pars opercularis (R) | 1700 |
|  | Pars orbitalis (R) | 1444 |
|  | Pars triangularis (R) | 1732 |
|  | Pericalcarine gyrus (R) | 716 |
|  | Postcentral gyrus (R) | 5160 |
|  | Posterior cingulate gyrus (R) | 3216 |
|  | Precentral gyrus (R) | 2228 |
|  | Precuneus (R) | 640 |
|  | Rostral anterior cingulate gyrus (R) | 1152 |
|  | Rostral middle frontal gyrus (R) | 2128 |
|  | Superior frontal gyrus (R) | 1696 |
|  | Superior parietal gyrus (R) | 1344 |
|  | Superior temporal gyrus (R) | 6824 |
|  | Supramarginal gyrus (R) | 1504 |
|  | Frontal pole (R) | 1732 |
|  | Temporal pole (R) | 756 |
|  | Transverse temporal gyrus (R) | 1048 |
|  | Insula (R) | 7812 |
|  | Thalamus (R) | 8324 |
|  | Caudate (R) | 2824 |
|  | Putamen (R) | 7140 |
|  | Pallidum (R) | 5272 |
|  | Hippocampus (R) | 1004 |
|  | Amygdala (R) | 872 |
|  | Accumbens (R) | 460 |

#### Table S8. Functional circuit differences between MDD and HC_test_

| Direction | Seed-based functional circuits | Δ percentage | *p* |
| --- | --- | --- | --- |
| Infra-normal |  |  |  |
|  | Fusiform gyrus (L) | − 10.37% | .039 |
|  | Lateral occipital gyrus (L) | − 10.98% | .019 |
|  | Pericalcarine gyrus (L) | − 7.32% | .042 |
|  | Rostral middle frontal gyrus (L) | − 7.32% | .042 |
|  | Superior temporal gyrus (L) | − 7.32% | .042 |
|  | Fusiform gyrus (R) | − 10.37% | .040 |
|  | Lateral occipital gyrus (R) | − 10.98% | .019 |
|  | Pericalcarine gyrus (R) | − 8.54% | .033 |
|  | Rostral middle frontal gyrus (R) | − 7.32% | .042 |
|  | Transverse temporal gyrus (R) | − 9.15% | .042 |
| Supra-normal |  |  |  |
|  | Fusiform gyrus (L) | 7.00% | .001 |
|  | Medial orbitofrontal gyrus (L) | 5.36% | < .001 |
|  | Precuneus (L) | 2.96% | .023 |
|  | Frontal pole (L) | 10.71% | < .001 |
|  | Bankssts (R) | 5.36% | < .001 |
|  | Fusiform gyrus (R) | 9.49% | < .001 |
|  | Lingual gyrus (R) | 7.67% | .010 |
|  | Precuneus (R) | 2.96% | .023 |
|  | Supramarginal gyrus (R) | 10.71% | < .001 |

Δ percentage is obtained by subtracting circuit percentage maps of HC_test_­ from circuit percentage map of MDD patients. The threshold of statistical significance was set at 0.05 and correction for multiple comparisons was performed using the false discovery rate (*p* < 0.05/3, FDR corrected). HC, healthy control; SZ, schizophrenia; L, left; R, right

#### Table S9. Functional circuit differences between BD and HC_test_

| Direction | Seed-based functional circuits | Δ percentage | *p* |
| --- | --- | --- | --- |
| Infra-normal |  |  |  |
|  | Cuneus (L) | 11.25% | .037 |
|  | Lateral occipital gyrus (L) | 23.11% | .001 |
|  | Pericalcarine gyrus (R) | 26.34% | < .001 |
|  | Supramarginal gyrus (R) | 11.86% | .004 |
| Supra-normal |  |  |  |
|  | Isthmus cingulate (L) | 9.68% | < .001 |
|  | Rostral anterior cingulate (L) | 6.45% | < .001 |
|  | Thalamus (L) | 6.45% | < .001 |
|  | Isthmus cingulate (R) | 9.68% | < .001 |
|  | Thalamus (R) | 6.45% | < .001 |
|  | Putamen (R) | 6.45% | < .001 |
|  | Pallidum (R) | 6.45% | < .001 |

Δ percentage is obtained by subtracting circuit percentage maps of HC_test_­ from circuit percentage map of BD patients. The threshold of statistical significance was set at 0.05 and correction for multiple comparisons was performed using the false discovery rate (*p* < 0.05/3, FDR corrected). HC, healthy control; SZ, schizophrenia; L, left; R, right

#### Table S10. Functional circuit differences between SZ and HC_test_

| Direction | Regions | Δ percentage | *p* |
| --- | --- | --- | --- |
| Infra-normal |  |  |  |
|  | Caudal middle frontal gyrus (L) | 10.35% | < .001 |
|  | Pars opercularis (L) | 6.85% | < .001 |
|  | Cuneus (R) | 10.65% | < .001 |
|  | Fusiform gyrus (R) | 3.50% | .011 |
|  | Inferior parietal gyrus (R) | 5.63% | < .001 |
|  | Pars opercularis (R) | 6.85% | < .001 |
|  | Pars triangularis (R) | 6.24% | .001 |
|  | Superior frontal gyrus (R) | 7.91% | .013 |
| Supra-normal |  |  |  |
|  | Bankssts (L) | 5.48% | < .001 |
|  | Cuneus (L) | 9.74% | < .001 |
|  | Fusiform gyrus (L) | 7.00% | .003 |
|  | Inferior temporal gyrus (L) | 4.87% | .005 |
|  | Lateral occipital gyrus (L) | 7.76% | .003 |
|  | Lingual gyrus (L) | 10.50% | < .001 |
|  | Pars opercularis (L) | 8.98% | < .001 |
|  | Pars orbitalis (L) | 4.11% | < .001 |
|  | Pars triangularis (L) | 6.24% | .002 |
|  | Pericalcarine gyrus (L) | 9.13% | .001 |
|  | Superior frontal gyrus (L) | 4.87% | .005 |
|  | Superior parietal gyrus (L) | 5.63% | .007 |
|  | Transverse temporal gyrus (L) | 4.87% | .004 |
|  | Bankssts (R) | 5.48% | < .001 |
|  | Caudal middle frontal gyrus (R) | 5.63% | .007 |
|  | Fusiform gyrus (R) | 8.37% | .001 |
|  | Inferior temporal gyrus (R) | 7.61% | .001 |
|  | Lateral occipital gyrus (R) | 7.76% | .003 |
|  | Lingual gyrus (R) | 7.76% | .003 |
|  | Pars opercularis (R) | 6.24% | .002 |
|  | Pars orbitalis (R) | 6.24% | .002 |
|  | Pars triangularis (R) | 6.24% | .002 |
|  | Pericalcarine gyrus (R) | 10.50% | < .001 |
|  | Posterior cingulate (R) | 6.24% | .001 |
|  | Precuneus (R) | 4.87% | .005 |
|  | Rostral middle frontal gyrus (R) | 5.63% | .007 |
|  | Superior parietal gyrus (R) | 5.63% | .007 |
|  | Transverse temporal gyrus (R) | 4.87% | .004 |

Δ percentage is obtained by subtracting circuit percentage maps of HC_test_­ from circuit percentage map of SZ patients. The threshold of statistical significance was set at 0.05 and correction for multiple comparisons was performed using the false discovery rate (*p* < 0.05/3, FDR corrected). HC, healthy control; SZ, schizophrenia; L, left; R, right

### Supplementary Figures

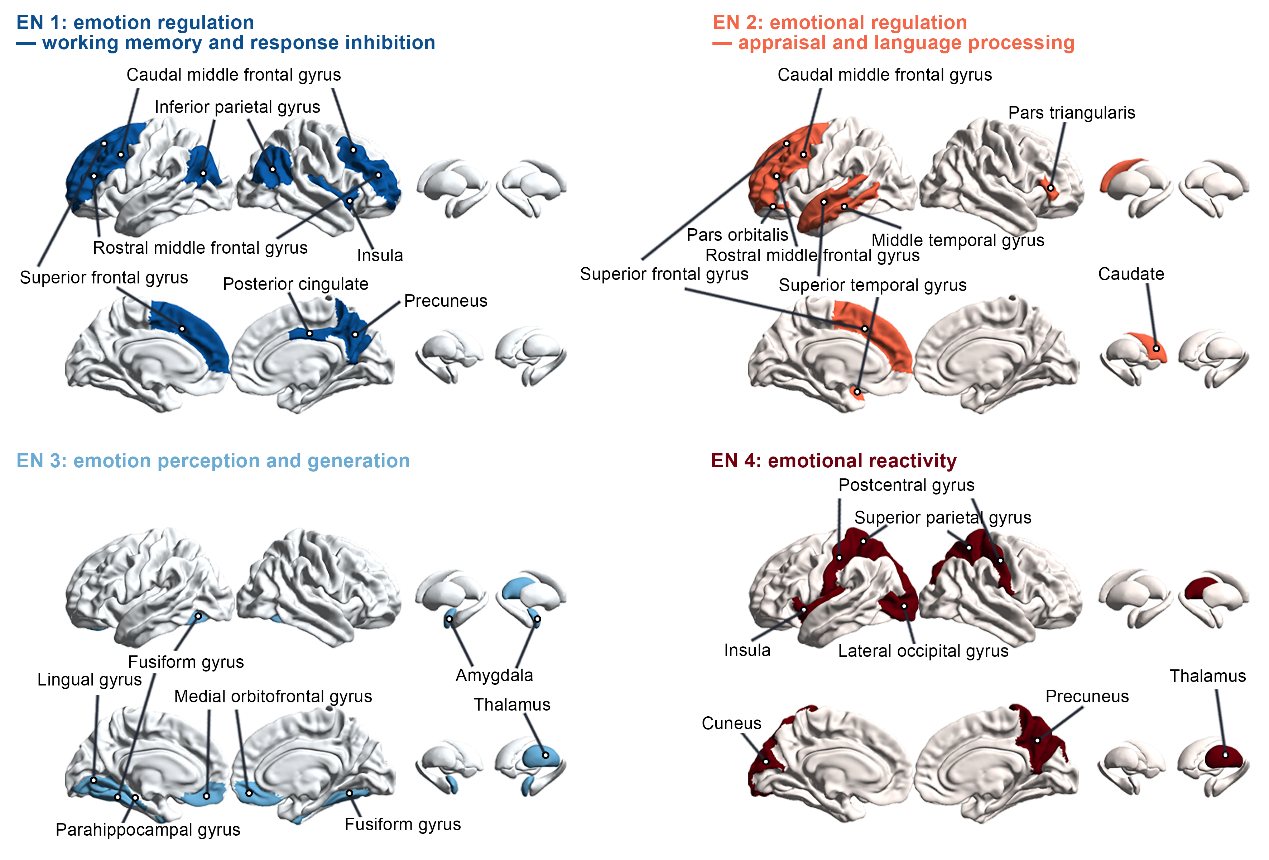

#### Figure S1. Large-scale emotional networks.

EN, emotional networks

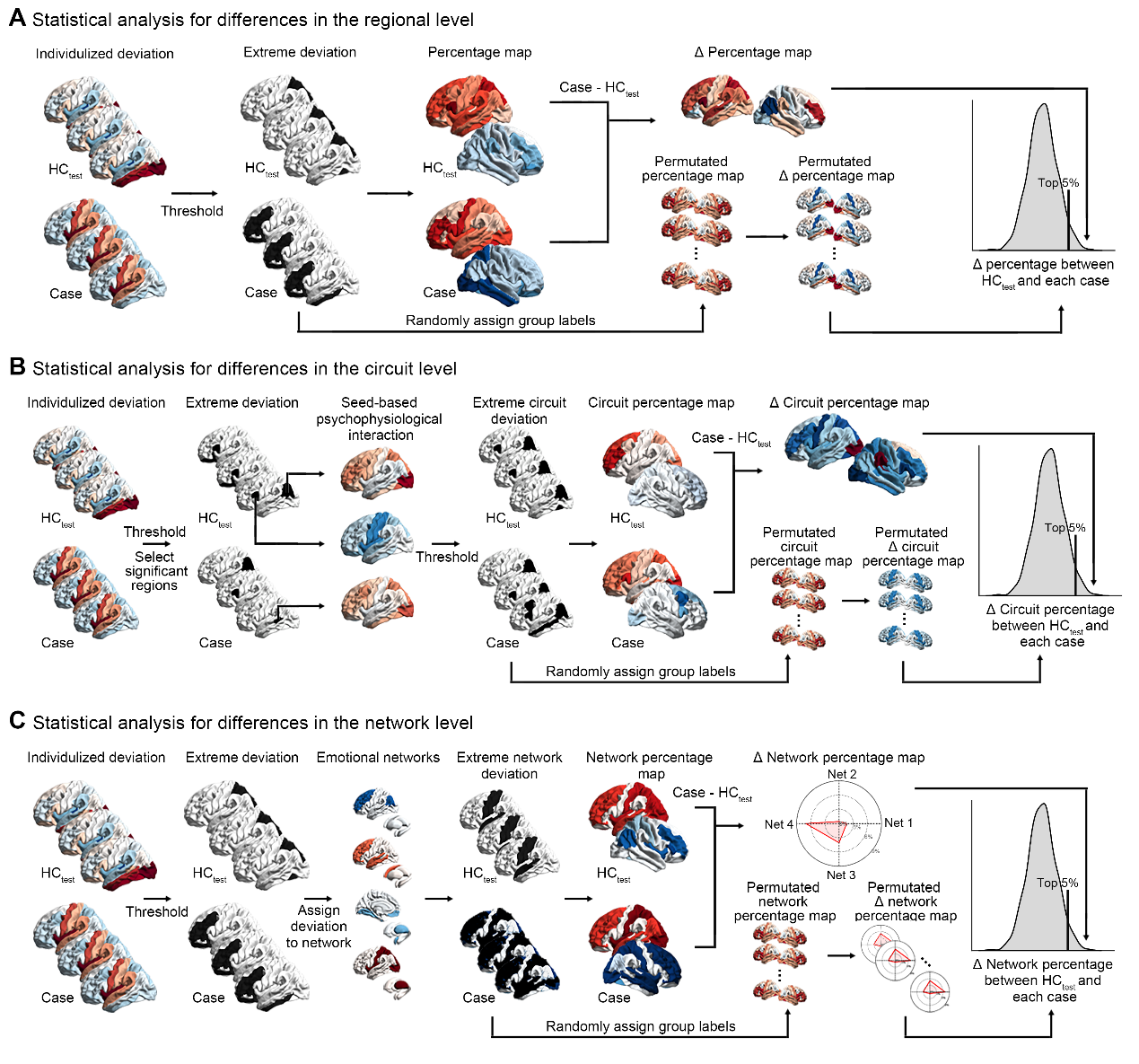

#### Figure S2. The pipeline of statistical analysis.

The pipeline of statistical analysis for differences in the regional **(A)**, circuit **(B)**, and network level **(C)**. HC, healthy controls

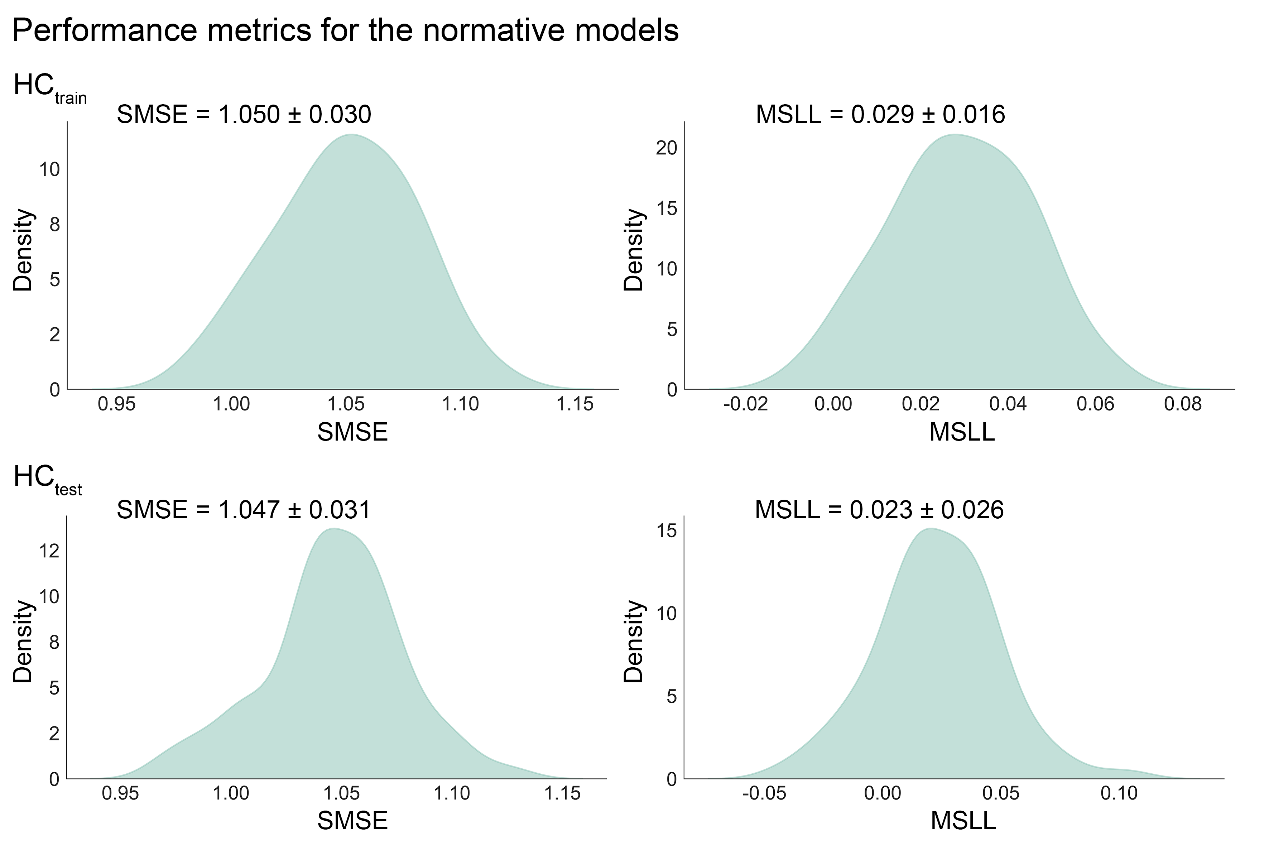

#### Figure S3. Performance metrics for the normative models of emotional episodic memory.

The distributions of SMSE and MSLL across emotional memory effect of all brain regions in HC_train_ (top) under 10-fold cross-validation, and HC_test_ (bottom) under a external validation. HC, healthy control; SMSE, standardized mean squared error; MSLL, mean squared log-loss

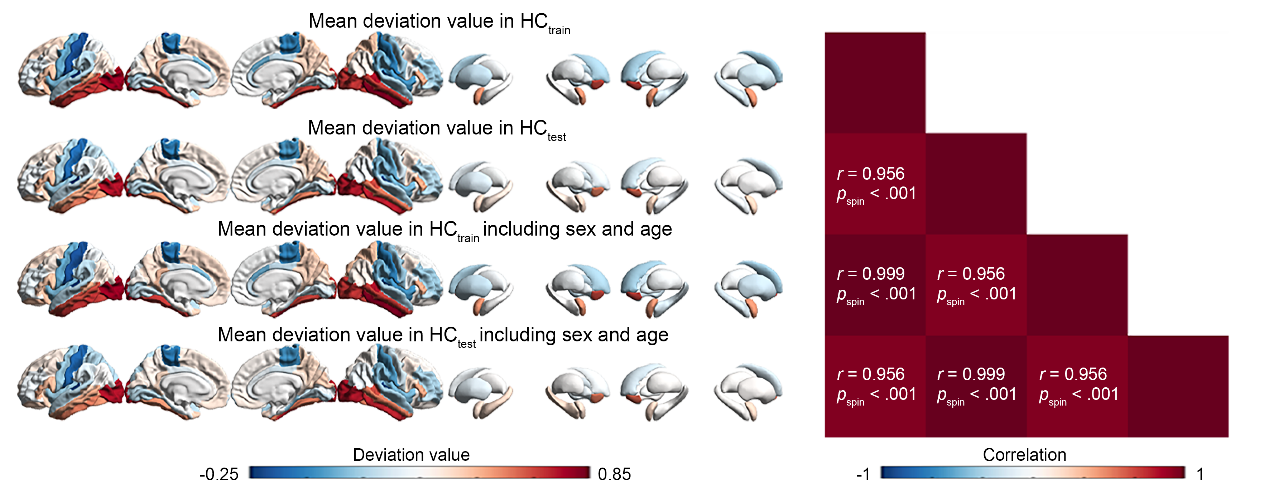

#### Figure S4. Age and sex effects on normative model.

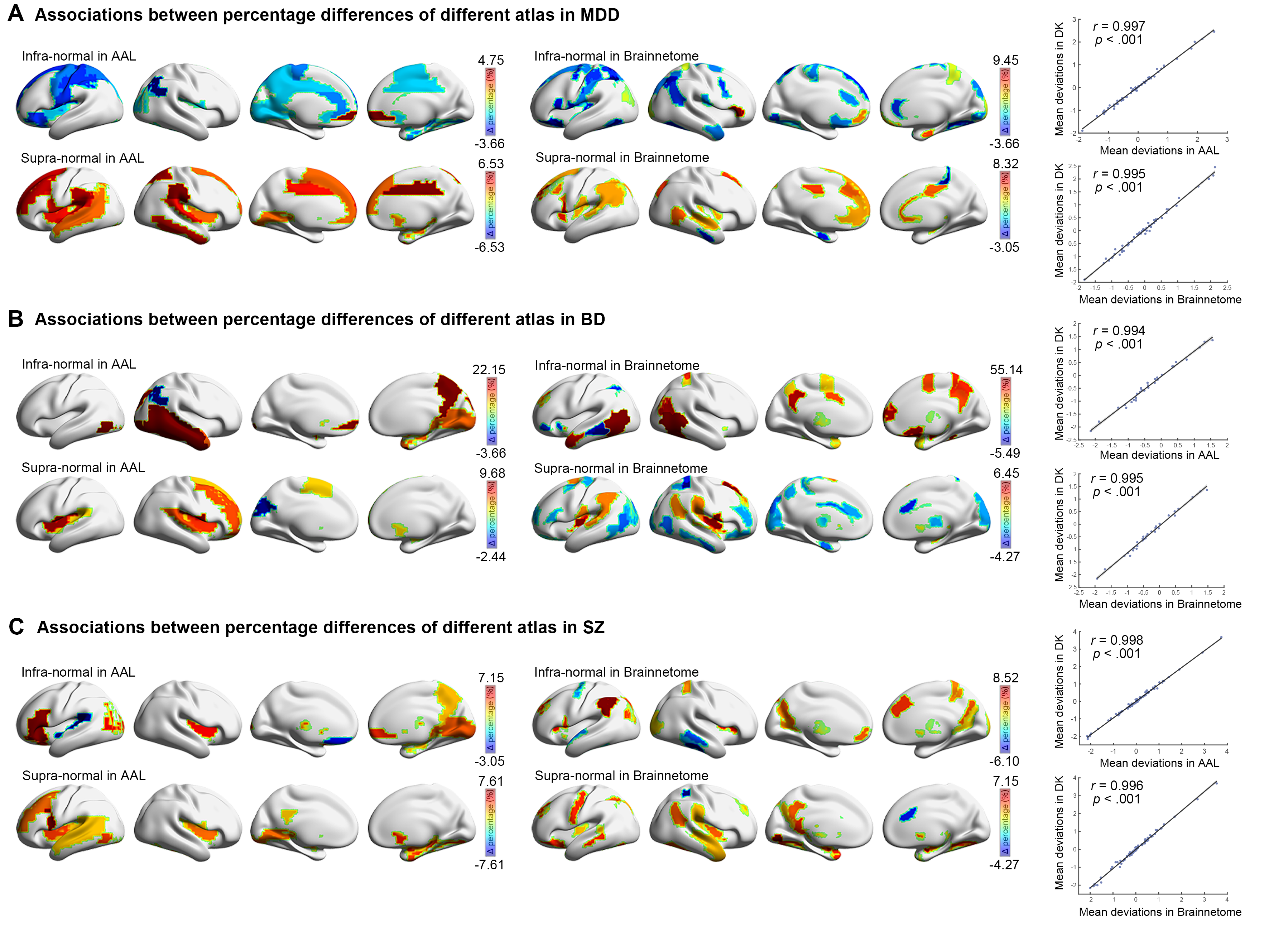

#### Figure S5. Association between percentage differences of different atlas.

The spatial distribution pattern of percentage differences exhibited high similarity across different brain parcellation atlas, and mean deviations in AAL and Brainnetome atlas were related to DK atlas in MDD **(A)**, BD **(B)** and SZ **(C)** patients. MDD, major depressive disorder; BD, bipolar disorder; SZ, schizophrenia; AAL, Anatomical Automatic Labelling; DK, Desikan-Killiany

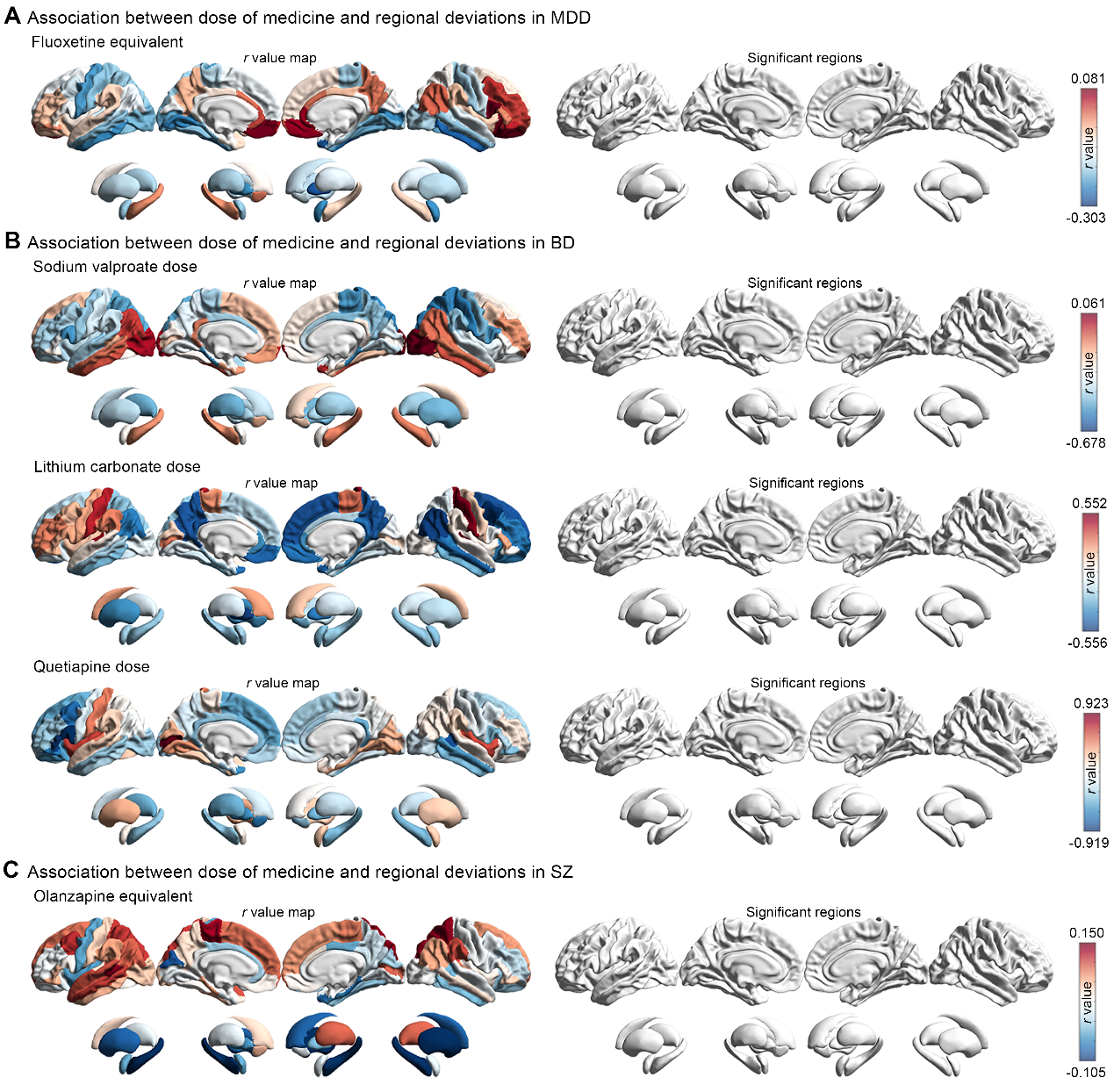

#### Figure S6. Association between dose of medicine and regional deviations.

No significant correlations between dose of medicine and regional deviations were be found in patients with MDD **(A)**, BD **(B)** and SZ **(C)**. The significant level was set at 0.05 and adjusted by FDR. MDD, major depressive disorder; BD, bipolar disorder; SZ, schizophrenia

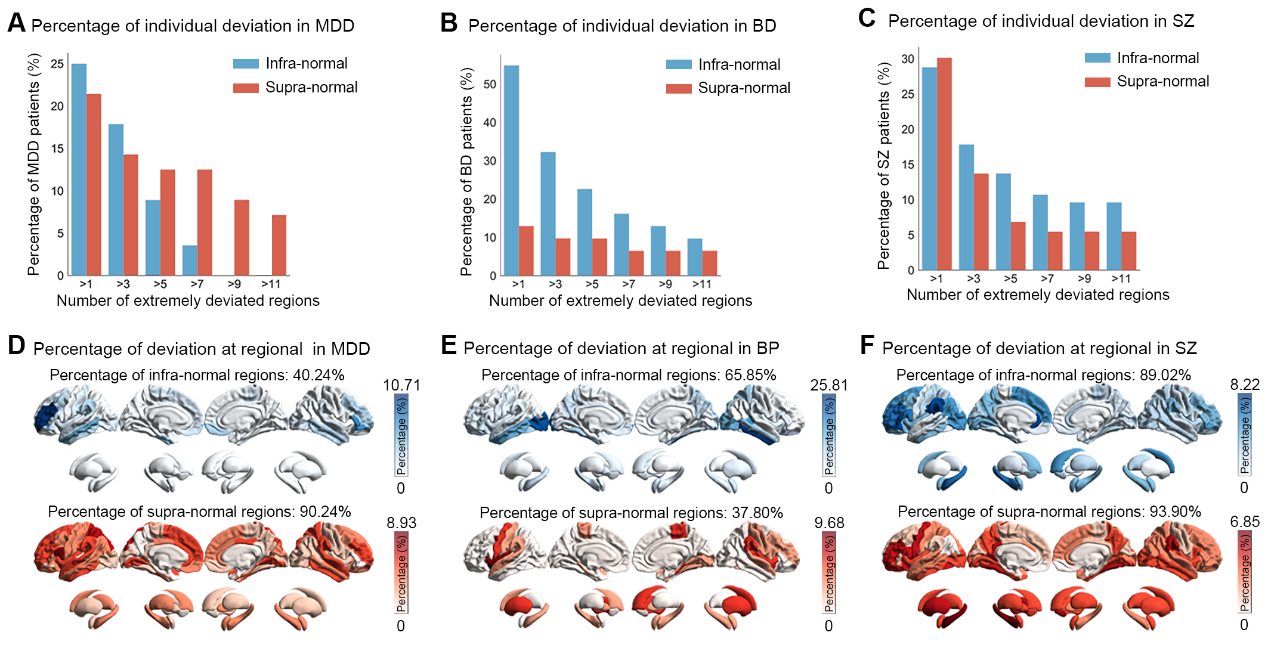

#### Figure S7. Regional heterogeneity of deviations of functional activation under “aversive vs. neutral” condition.

Bar plots showed the distribution of the number of regions per patient with extremely supra-normal (red) and infra-normal (blue) deviations in MDD **(A)**, BD **(B)**, and SZ patients **(C)**. The spatial overlap maps indicated the percentage of patients who deviated extremely from the normative range for each brain region in MDD **(D)**, BD **(E)**, and SZ patients **(F)**. MDD, major depressive disorder; BD, bipolar disorder; SZ, schizophrenia

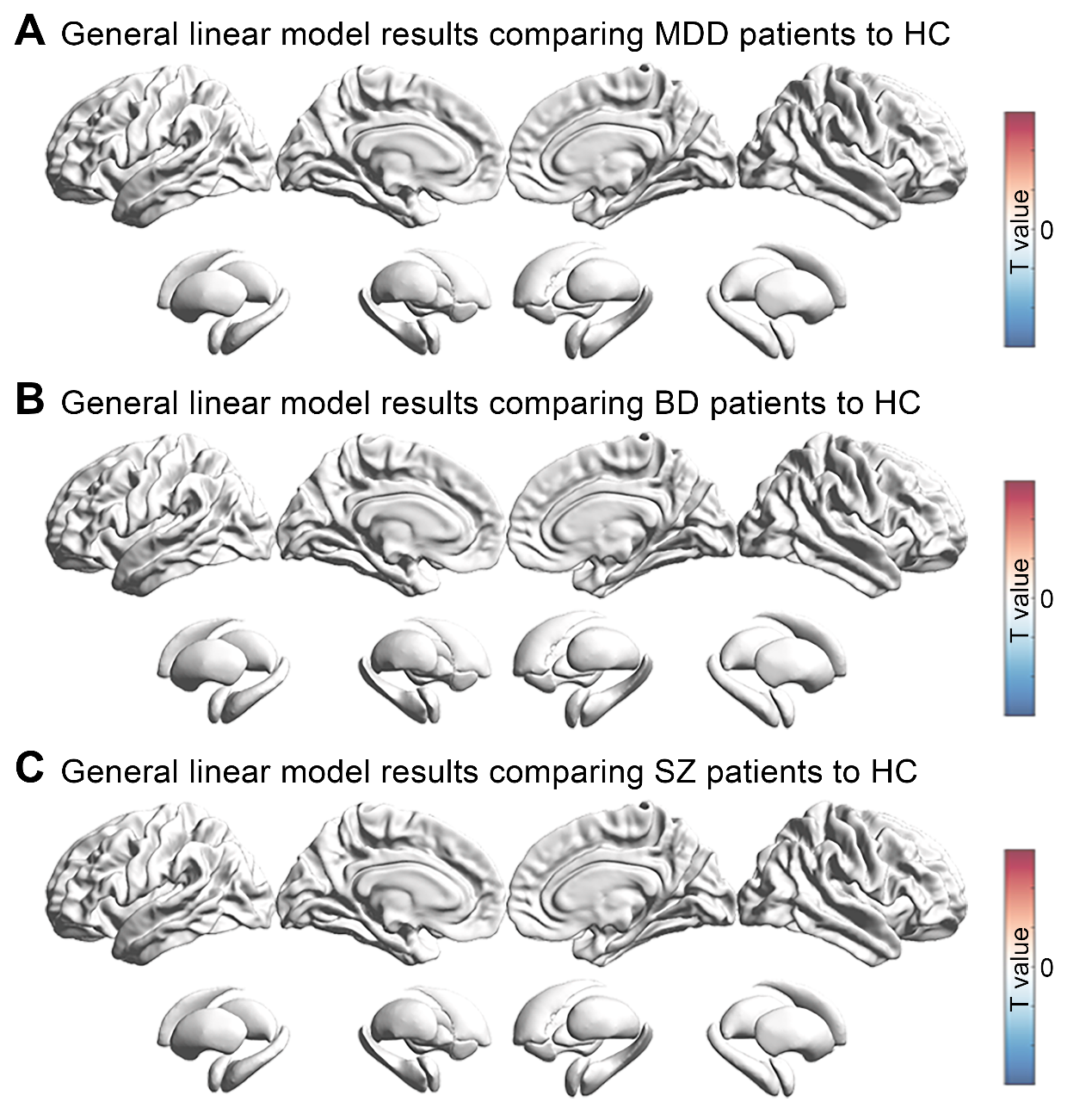

#### Figure S8. Traditional group-level comparison using general linear model.

There are no significant regions in general linear model comparing MDD **(A)**, BD **(B)** and SZ **(C)** patients to HC, respectively. MDD, major depressive disorder; BD, bipolar disorder; SZ, schizophrenia; HC, healthy control

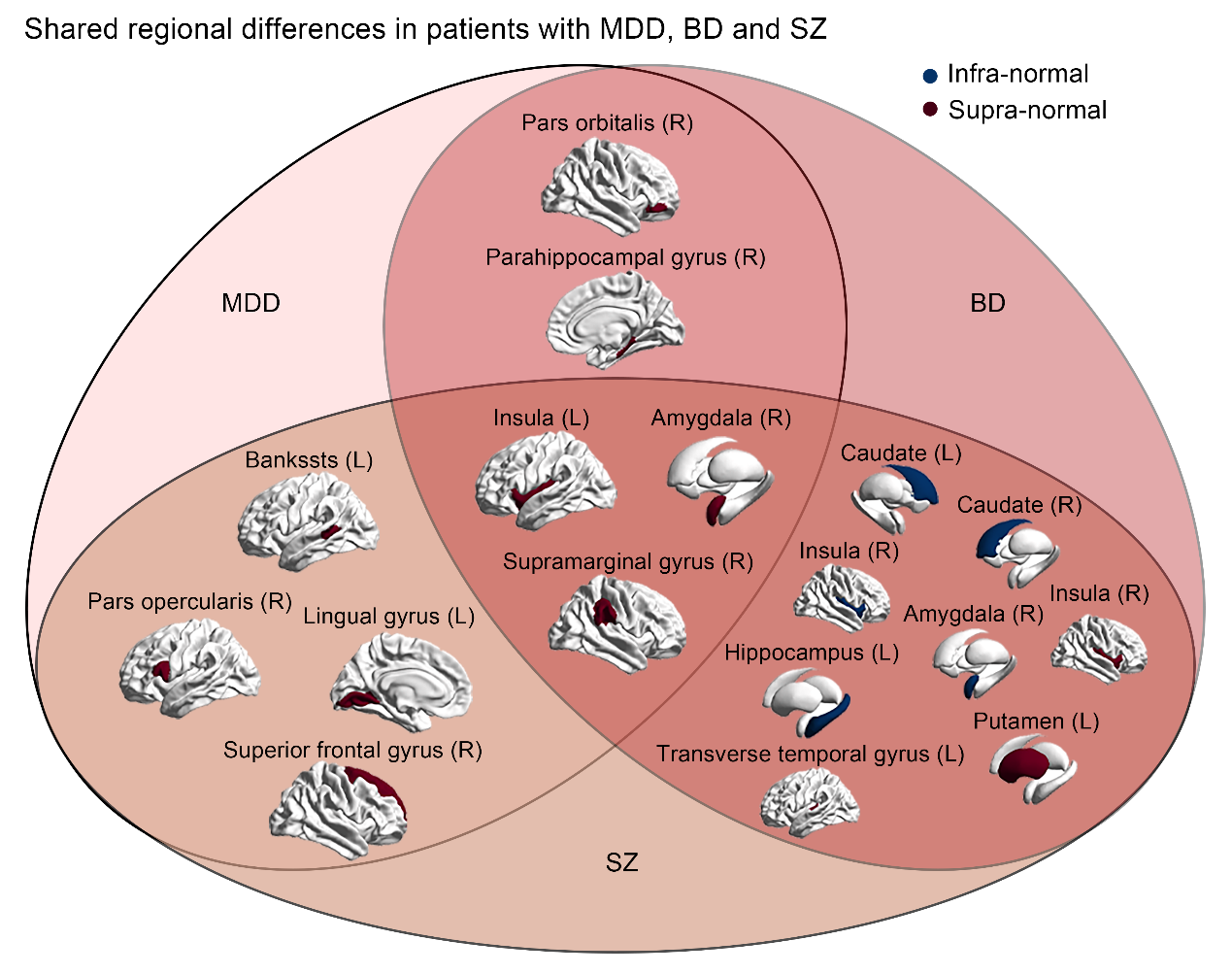

#### Figure S9. The shared regional differences in patients with MDD, BD and SZ.

Shared map of regional differences was constructed by integrating brain regions with extreme deviation shared among any two or three psychiatric disorders. MDD, major depressive disorder; BD, bipolar disorder; SZ, schizophrenia

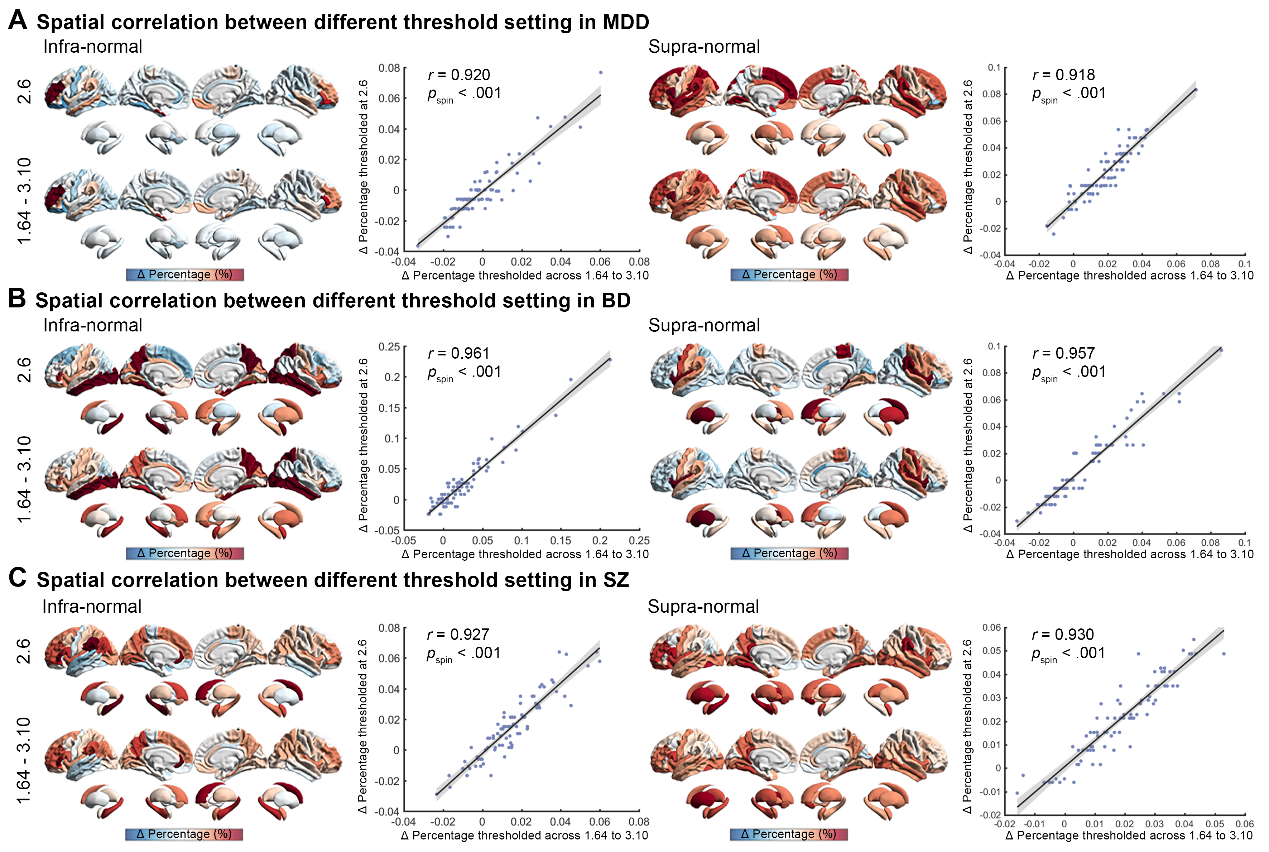

#### Figure S10. Spatial correlation between different threshold setting in percentage differences map of all case groups.

There are significant spatial correlations in infra-normal and supranormal regions of MDD **(A)**, BD **(B)** and SZ **(C)** patients. The significant level was set at 0.05 with 10,000 times permutation tests with spatial autocorrelation and adjusted by FDR. MDD, major depressive disorder; BD, bipolar disorder; SZ, schizophrenia

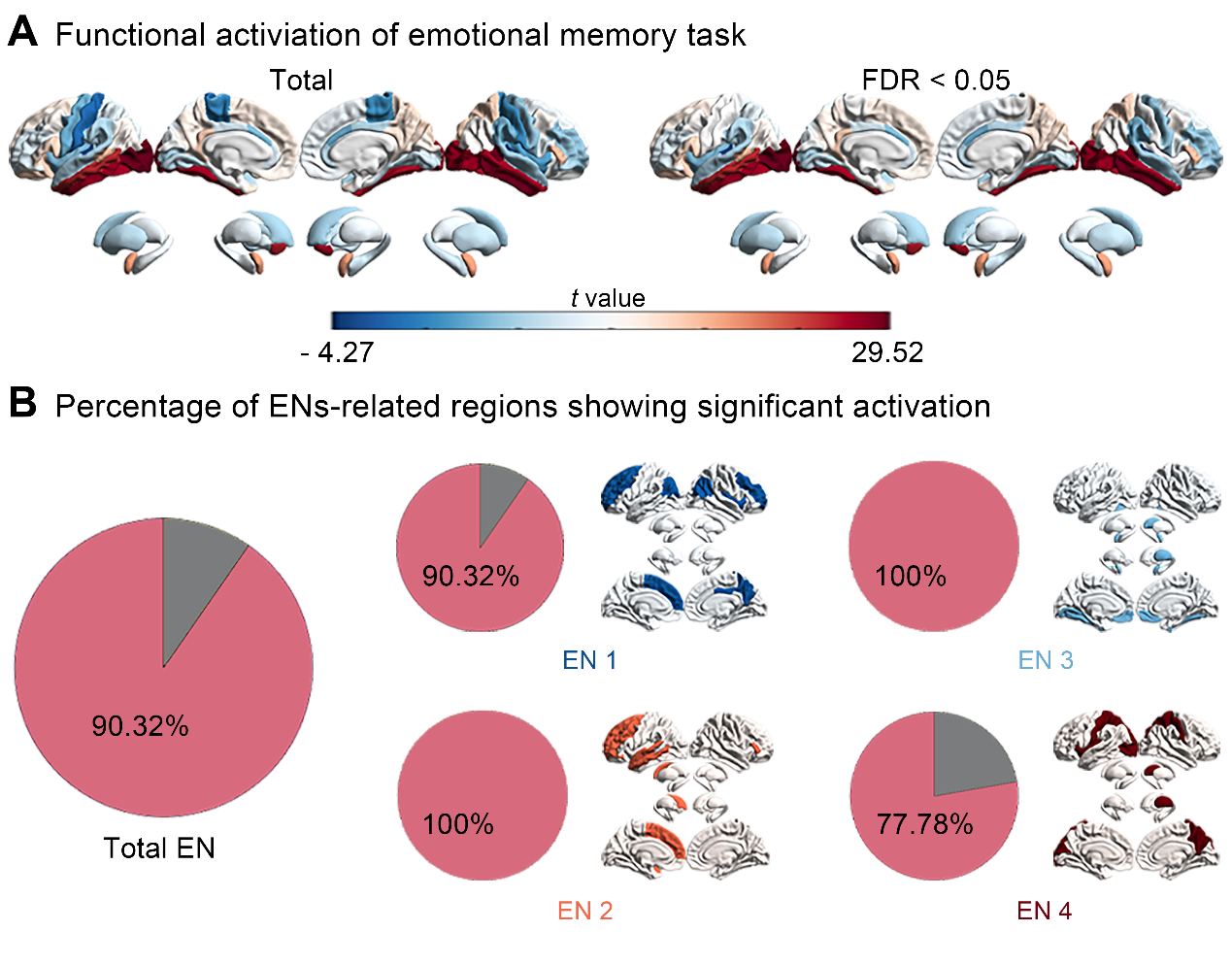

#### Figure S11. Association between functional activation at “aversive vs. neutral” condition and emotional networks

**(A)** Significant functional activation at “aversive vs. neutral” condition was evaluated using one-sample *t*-test. **(B)** The regions with significant activation were embedded at four emotional networks, showing high percentage of ENs-related regions showing signification activation. FDR, false discovery rate; EN, emotional network

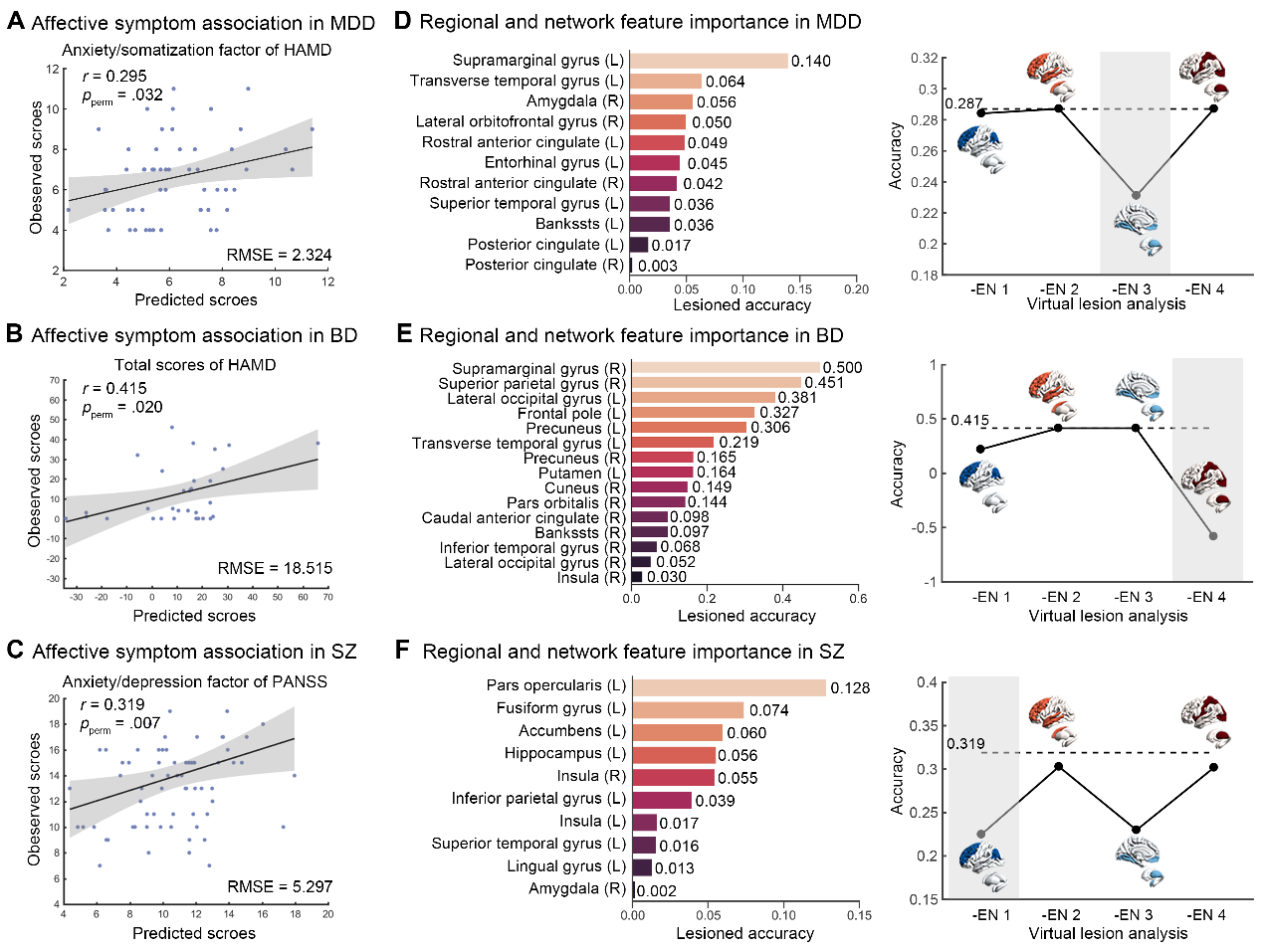

#### Figure S12. The important features in predicted model of affective model.

The scatter plot showed the prediction accuracy defined by correlation coefficient between observed scores and predictive scores. **(A)** Individual deviations significantly predicted the anxiety/somatization factor of HAMD in MDD patients. **(B)** Total scores of HAMD in BD patients were predicted based on individual deviations. **(C)** Individual deviations predicted the anxiety/depression factor of PANSS in SZ patients. The virtual lesion analysis was applied to identify important features in prediction model of MDD **(D)**, BD **(E)** and SZ patients **(F)**. The bar plot exhibited the regional predictive weights (middle) and line chart showed the predictive weights at network level (right). In line chart, the dotted line indicates the original prediction accuracy and the network that exhibits the largest feature weight marked by shadow. MDD, major depressive disorder; BD, bipolar disorder; SZ, schizophrenia; RMSE, root mean square error; HAMD, Hamilton depression scale; PANSS, positive and negative syndrome scale; EN, emotional network; R, right; L, left

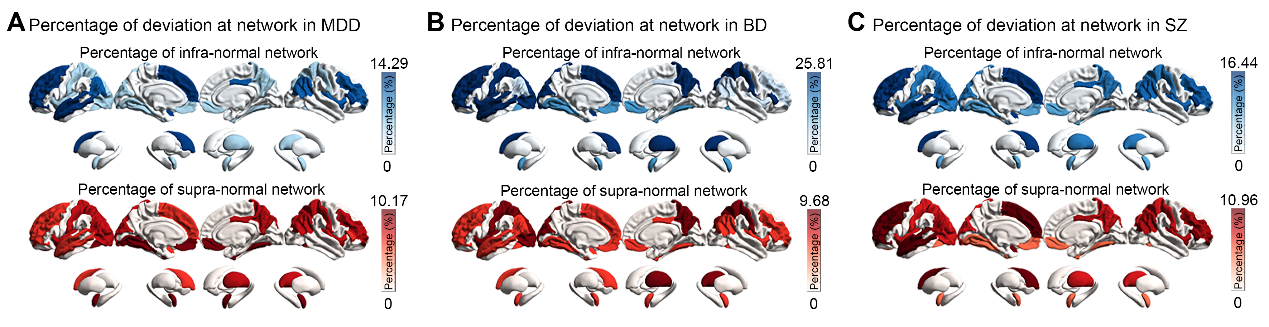

#### Figure S13. Network heterogeneity of deviation of functional activation under “aversive vs. neutral” condition.

The spatial overlap maps indicated the percentage of patients who deviated extremely from the normative range for each emotional network in MDD **(A)**, BD **(B)**, and SZ patients **(C)**. MDD, major depressive disorder; BD, bipolar disorder; SZ, schizophrenia

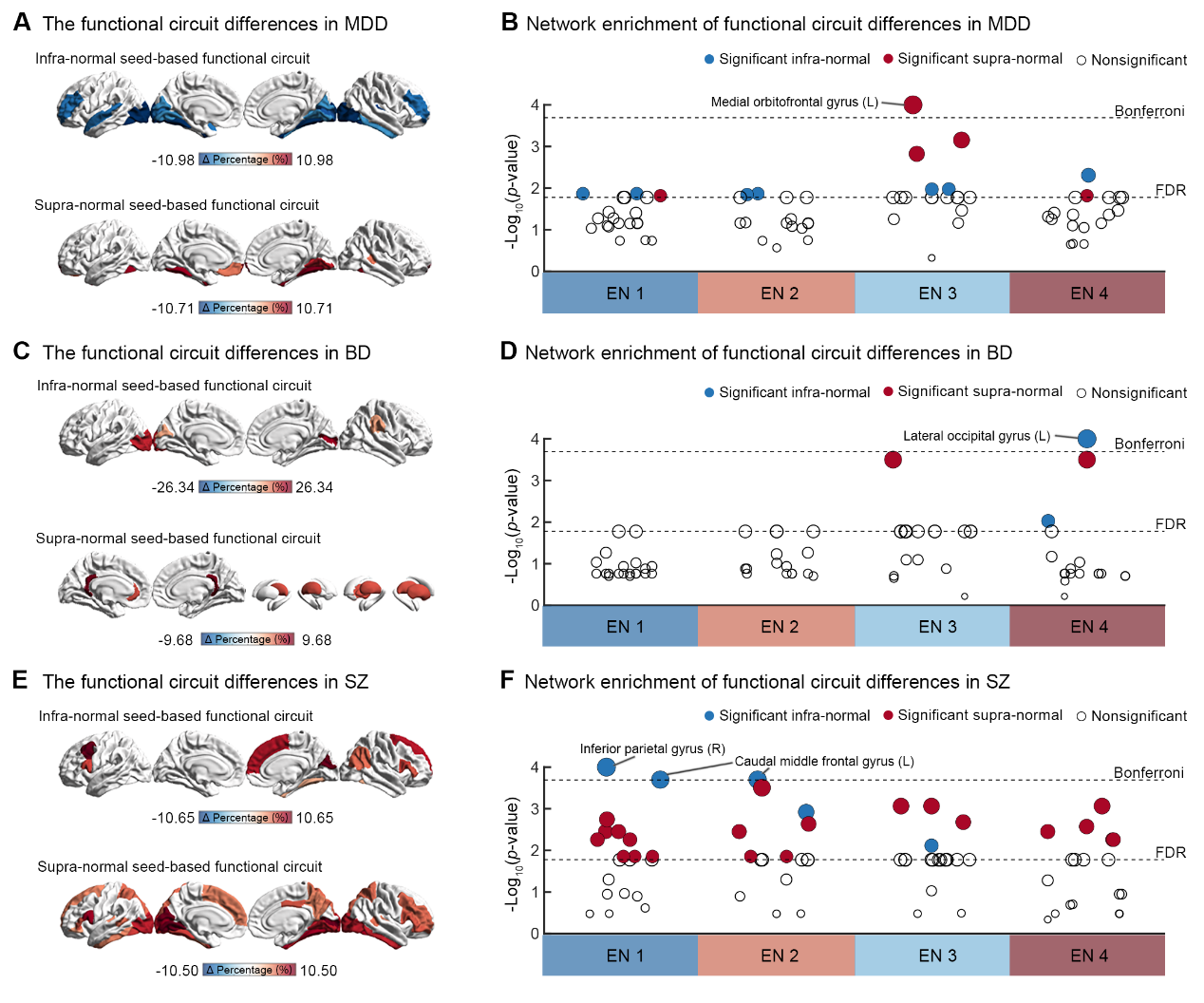

#### Figure S14. Group convergent effect of individual deviations at circuit.

The group convergent effect of individual deviations at circuit level based on seeds showing significant deviations in MDD **(A)**, BD **(C)** and SZ **(E)** patients were shown. Significant circuit deviations were further located into four ENs for MDD **(D)**, BD **(B)** and SZ **(F)** patients, with blue and red colors respectively indicated the infra-normal and supra-normal deviations and circle size proportional to the significant level. The statistically significance was determined by using 10,000 times group-base permutation test and adjusted by FDR. The most significant differences (Bonferroni correction) were labeled. MDD, major depressive disorder; BD, bipolar disorders; SZ, schizophrenia; EN, emotional network

### References

1.	Acton PD, Friston KJ (1998): Statistical parametric mapping in functional neuroimaging: beyond PET and fMRI activation studies. *European journal of nuclear medicine*. 25:663-667.

2.	Chen Q, Ursini G, Romer AL, Knodt AR, Mezeivtch K, Xiao E, et al. (2018): Schizophrenia polygenic risk score predicts mnemonic hippocampal activity. *Brain*. 141:1218-1228.

3.	Lang PJ, Bradley MM, Cuthbert BN (1998): Emotion and motivation: measuring affective perception. *Journal of clinical neurophysiology*. 15:397-408.

4.	Lang PJ, Bradley MM, Cuthbert BN (1990): Emotion, attention, and the startle reflex. *Psychological review*. 97:377-395.

5.	Arnatkeviciute A, Fulcher BD, Fornito A (2019): A practical guide to linking brain-wide gene expression and neuroimaging data. *NeuroImage*. 189:353-367.

6.	Morawetz C, Riedel MC, Salo T, Berboth S, Eickhoff SB, Laird AR, et al. (2020): Multiple large-scale neural networks underlying emotion regulation. *Neuroscience and biobehavioral reviews*. 116:382-395.

7.	Boes AD, Prasad S, Liu H, Liu Q, Pascual-Leone A, Caviness VS, Jr., et al. (2015): Network localization of neurological symptoms from focal brain lesions. *Brain*. 138:3061-3075.

8.	Fox MD (2018): Mapping symptoms to brain networks with the human connectome. *The New England journal of medicine*. 379:2237-2245.

9.	Segal A, Parkes L, Aquino K, Kia SM, Wolfers T, Franke B, et al. (2023): Regional, circuit and network heterogeneity of brain abnormalities in psychiatric disorders. *Nature neuroscience*. 26:1613-1629.

10.	Borne L, Tian Y, Lupton MK, van der Meer JN, Jeganathan J, Paton B, et al. (2023): Functional re-organization of hippocampal-cortical gradients during naturalistic memory processes. *NeuroImage*. 271:119996.

11.	Tian Y, Margulies DS, Breakspear M, Zalesky A (2020): Topographic organization of the human subcortex unveiled with functional connectivity gradients. *Nature neuroscience*. 23:1421-1432.

12.	Desikan RS, Ségonne F, Fischl B, Quinn BT, Dickerson BC, Blacker D, et al. (2006): An automated labeling system for subdividing the human cerebral cortex on MRI scans into gyral based regions of interest. *NeuroImage*. 31:968-980.

13.	Fastenrath M, Spalek K, Coynel D, Loos E, Milnik A, Egli T, et al. (2022): Human cerebellum and corticocerebellar connections involved in emotional memory enhancement. *Proceedings of the National Academy of Sciences of the United States of America*. 119:e2204900119.

14.	Friston KJ, Buechel C, Fink GR, Morris J, Rolls E, Dolan RJ (1997): Psychophysiological and modulatory interactions in neuroimaging. *NeuroImage*. 6:218-229.

15.	Ongür D, Price JL (2000): The organization of networks within the orbital and medial prefrontal cortex of rats, monkeys and humans. *Cerebral cortex*. 10:206-219.

16.	Shiba Y, Oikonomidis L, Sawiak S, Fryer TD, Hong YT, Cockcroft G, et al. (2017): Converging prefronto-insula-amygdala pathways in negative emotion regulation in marmoset monkeys. *Biological psychiatry*. 82:895-903.

17.	Gottfried JA, Dolan RJ (2004): Human orbitofrontal cortex mediates extinction learning while accessing conditioned representations of value. *Nature neuroscience*. 7:1144-1152.

18.	Bo K, Kraynak TE, Kwon M, Sun M, Gianaros PJ, Wager TD (2024): A systems identification approach using Bayes factors to deconstruct the brain bases of emotion regulation. *Nature neuroscience*. 27:975-987.

19.	Guimond S, Padani S, Lutz O, Eack S, Thermenos H, Keshavan M (2018): Impaired regulation of emotional distractors during working memory load in schizophrenia. *Journal of psychiatric research*. 101:14-20.
